# Enhanced STAT5a activation rewires exhausted CD8 T cells during chronic stimulation to acquire a hybrid durable effector like state

**DOI:** 10.1101/2022.10.03.509766

**Authors:** Jean-Christophe Beltra, Mohamed S. Abdel-Hakeem, Sasikanth Manne, Zhen Zhang, Hua Huang, Makoto Kurachi, Leon Su, Lora Picton, Yuki Muroyama, Valentina Casella, Yinghui J. Huang, Josephine R. Giles, Divij Mathew, Jonathan Belman, Max Klapholz, Hélène Decaluwe, Alexander C. Huang, Shelley L. Berger, K. Christopher Garcia, E. John Wherry

## Abstract

Rewiring exhausted CD8 T cells (T_EX_) towards more functional states is a major goal of cancer immunotherapy but has proven challenging due to the epigenetic stability of T_EX_. Indeed, T_EX_ are epigenetically programmed by the transcription factor Tox. However, epigenetic changes continue to occur as T_EX_ transition from progenitor (T_EX_^prog^), to intermediate (T_EX_^int^) and terminal (T_EX_^term^) subsets, suggesting potential developmental flexibility in mature T_EX_ subsets. By examining the transition of T_EX_^prog^ into T_EX_^int^ cells, we discovered a reciprocally antagonistic circuit between Stat5a and Tox in T_EX_ cells. Stat5-activity controlled T_EX_^int^ development, antagonized Tox, and instigated partial effector biology. Stat5 was also essential for T_EX_ reinvigoration by PD-1 blockade. Indeed, temporal induction of Stat5-activity in T_EX_ using an orthogonal IL-2/IL2Rβ-pair fostered T_EX_^int^ cell accumulation and synergized with PD-L1 blockade. Constitutive Stat5a activity (STAT5CA) antagonized Tox-dependent T_EX_ epigenetic programming to generate a durable hybrid effector/NK-like population with enhanced tumor control. Finally, enforcing Stat5-signals in established T_EX_^prog^ partially rewired the T_EX_ epigenetic landscape towards the effector/memory lineage. Together, these data highlight therapeutic opportunities of manipulating Stat5 to rewire T_EX_ towards a durably protective hybrid program.

## INTRODUCTION

CD8 T cell exhaustion is a common feature of chronic viral infections and cancers and limits effective control of disease^1, 2^. T cell exhaustion is also a major barrier to effective immunotherapy for cancer including engineered cellular therapies^3^ and has been implicated in autoimmune diseases.^2^ First described as loss of effector functions in settings of chronic antigenic stimulation,^4, 5^ it is now clear that CD8 T cell exhaustion is a distinct epigenetically programmed state of T cell differentiation initiated by the HMG-transcription factor (TF) Tox.^6–10^ This Tox-dependent epigenetic program also precludes (re)differentiation of T_EX_ towards functional effector (T_EFF_) or memory (T_MEM_) CD8 T cells.^11–14^ Despite this altered differentiation state, T_EX_ acquire the unique ability to persist long-term despite chronic antigenic stimulation,^14, 15^ and exert partial control of chronic infections and cancer.^1, 16–18^ Moreover, relieving some inhibition of T_EX_ through PD-1 pathway blockade results in considerable clinical efficacy for some cancers.^19–24^ Nevertheless, PD-1 pathway blockade fails to induce permanent epigenetic rewiring of T_EX_^25, 26^ highlighting the need to identify strategies that can relieve the constraints on (re)differentiation of T_EX_ cells and allow conversion to more durably functional CD8 T cell states.

Recent work examining the developmental biology of T_EX_ cells has defined biologically distinct subsets that are related in a developmental hierarchy.^27–30^ Molecular characterization of T_EX_ subsets has provided insights into subset-specific biology and their therapeutic relevance. For example, a Tcf1 (*Tcf7*)-expressing progenitor (T_EX_^prog^) subset retains proliferative potential and generates downstream T_EX_ subsets in the steady state.^27, 31–33^ These T_EX_^prog^ exist in two interchangeable states including a stem-like - quiescent subset (T_EX_^prog1^) that resides in lymphoid tissues and a transcriptionally distinct subpopulation (T_EX_^prog2^) that leaves these lymphoid niches and re-enters cell cycle. As these T_EX_^prog2^ cells proliferate, they lose Tcf1 expression and differentiate into an intermediate “effector-like” subset (T_EX_^int^) that circulates in blood and ultimately converts to terminally exhausted CD8 T cells (T_EX_^term^) upon entering peripheral tissues where these T_EX_^term^ also acquire features of tissue residency.^27^ An additional set of insights involves the T_EX_^int^ subset. First, PD-1 blockade functions as a transient amplifier of this T_EX_^int^ population.^27, 28^ Although T_EX_^prog^ initiate the proliferative response to PD-1 blockade,^31, 32^ subsequent production of new “effector-like” T_EX_^int^ cells likely mediates the therapeutic benefits. Second, partial re-acquisition of effector features by T_EX_^int^ cells in the landscape of a Tox-dependent exhaustion program suggests possible developmental flexibility. The molecular mechanisms of this potential T_EX_ rewiring, however, remain poorly understood. Thus, identifying mechanisms controlling T_EX_ subset conversions and biology, particularly at the T_EX_^int^ stage could reveal opportunities to rewire the differentiation trajectory of T_EX_ for therapeutic benefit.

Here, we discovered a reciprocal antagonistic circuit between Tox and Stat5a in T_EX_ including preferential re-engagement of Stat5a activity in the T_EX_^int^ subset. Indeed, Stat5 was essential for T_EX_^int^ cell development as well as for the response to PD-1 pathway blockade. Moreover, Stat5 controlled re-activation of effector-like machinery acquired at the T_EX_^int^ stage, fostering cytolytic effector functions even in the context of a T_EX_ chromatin landscape. Enforcing constitutive Stat5a activity (STAT5CA) antagonized Tox and the Tox-dependent T_EX_ programing in chronic viral infection. This enforced Stat5a activity resulted in the induction of a suite of effector and NK-like biology in virus-specific CD8 T cells. These changes were driven by re-wired epigenetic and transcriptional circuitry and resulted in durable accumulation, and superior protective capacity, of STAT5CA-expressing CD8 T cells in settings of chronic viral infection and cancer. Therapeutic delivery of IL-2/Stat5-signals selectively to T_EX_ cells using an orthogonal IL-2/IL2Rβ pair system enhanced formation of T_EX_^int^ cells and synergized robustly with PD-1 blockade. Finally, temporal reactivation of Stat5 in T_EX_^prog^ reversed key epigenetic features of exhaustion, restored accessibility at T_EFF_/T_MEM_-related open chromatin regions and improved polyfunctionality. Together, these observations identify Stat5 as a key regulator of T_EX_ differentiation, antagonizing Tox-driven terminal exhaustion and fostering improved effector activity and durability in the setting of chronic antigen (Ag) stimulation. Moreover, accessing this biology by therapeutic augmenting Stat5 signals specifically in T_EX_ in combination with PD-1 blockade not only expanded the T_EX_^int^ population but also rewired these T_EX_ cells towards a more protective differentiation state with features of durability under chronic antigenic stress and enhanced effector biology.

## Results

### Tox restrains Stat5a activity in virus-specific CD8 T cells during chronic infection

During chronic viral infections and cancer, Tox fosters epigenetic commitment of Ag-specific CD8 T cells towards exhaustion.^6–9^ This fate commitment to T_EX_ is mediated in part by antagonizing pathways of functional effector (T_EFF_) and memory (T_MEM_) cell differentiation. However, partial transcriptional and epigenetic re-engagement of effector biology occurs at the T_EX_^int^ cell stage suggesting the existence of yet unidentified pro-effector molecular circuits capable of temporally counterbalancing the Tox-dependent exhaustion program.^27^ To identify potential mechanisms of re-engagement of effector circuitry in T_EX_ cells, we investigated transcriptional circuits and upstream regulators preferentially antagonized by Tox in virus-specific CD8 T cells during chronic viral infection. We performed Ingenuity <u>P</u>athways <u>A</u>nalysis (IPA) on differentially expressed genes (DEGs) between WT and Tox-deficient TCR transgenic “P14” CD8 T cells specific for LCMV D^b^gp_33-41_ from previously published RNA sequencing (RNAseq) data (**Fig. S1A**).^7^ Networks of transcriptional regulators involved in interferon signaling (i.e. *Irf1*, *Irf7*, *Stat1*) and terminal exhaustion (i.e. *Foxo1*) enriched in WT P14 cells. Conversely, TFs associated with T_EFF_ or T_MEM_ differentiation (i.e. *Tbx21*, *Id3* and *Stat5a*) had elevated transcriptional activity in Tox-deficient P14 cells (**Fig. S1B, Table S1**). Using a second single cell RNA sequencing (scRNAseq) dataset,^6^ transcriptional networks of Tbet and Stat5a again enriched in Tox-deficient compared to WT D^b^gp33^+^ CD8 T cells and these TFs scored among the top 10 regulators enriched in 3 out of the 5 main clusters of cells identified (**Fig. S1C-G, Table S1**). Among the transcriptional regulators identified by IPA, Tbet and Stat5a overlapped with a list of TFs independently predicted using Taiji analysis^34^ to selectively impact T_EX_^int^ cells compared to other T_EX_ subsets (**Fig. S1H**).^27^ Previous work identified bi-directional antagonism between Tbet and Tox in T_EX_^int^ cells.^27^ However, Taiji analysis predicted Stat5a activity to be even more specifically enriched in the T_EX_^int^ subset than Tbet, and the IPA-defined transcriptional network of Stat5a was more strongly anti-correlated with Tox expression than Tbet (**Fig. 1A and S1H**). These analyses suggested a possible antagonistic axis between Tox and Stat5a in T_EX_ cells. Indeed, the transcriptional signatures of Stat5a and Tox were inversely correlated in Ag-specific CD8 T cells at 1 week and 1 month of chronic infection (**Fig. 1B-C**). Stat5a activity in T_EX_ cells was also anti-correlated with the expression of exhaustion-specific genes, but positively associated with genes involved in effector-related biology (**Fig. S1I**). These data suggested a role for Stat5a in antagonizing Tox and the program of CD8 T cell exhaustion.

**Figure 1:**
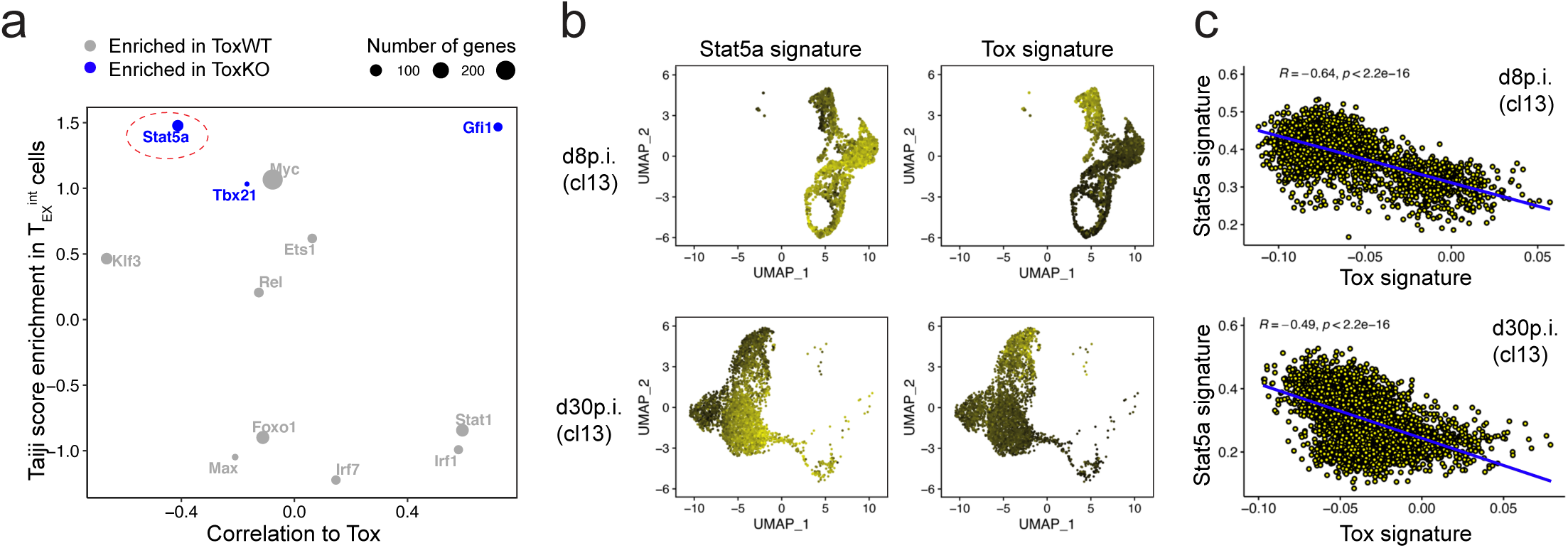
Reciprocal Activity between Stat5a and Tox in Ag-specific CD8 T cells during chronic infection. **A**-Dot plot of IPA regulators significantly enriched from **Fig. S1B** (logp≥10) that were also enriched in an independent Taiji Rank analysis in **Fig. S1H**. Selected IPA regulators are plotted based on their fold change in Taiji enrichment for T_EX_^int^ cells (Y axis) and correlation of the IPA-defined gene network for each TF to Tox expression (X-axis). **B**-UMAP of re-processed scRNAseq of P14 CD8 T cells at d8 (upper panel)^35^ or d30 (lower panel)^11^ post Cl13 infection projecting the Stat5a network as defined by the IPA analysis (left) or a core signature for Tox (genes enriched in P14 WT versus P14 ToxKO).^7^ **C**-Correlation scores between Stat5a network and a Tox signature at indicated time of Cl13 infection.

### STAT5a reduces Tox expression, antagonizes exhaustion and fosters effector-like CD8 T cell differentiation during chronic viral infection

During the first week of a developing chronic viral infection, Ag-specific CD8 T cells differentiate into either effector-like CD8 T cells or precursors of T_EX_.^26, 35^ Because the potential Stat5a and Tox antagonism observed above was evident by d8p.i., we asked whether Stat5a could impact early CD8 T cell-fate commitment during chronic infection. Congenically distinct P14 cells were transduced with retroviruses (RV) encoding either a constitutively active form of Stat5a (P14 STAT5CA)^36, 37^ or a control RV (P14 Empty). RV-transduced P14 cells were sort-purified based on RV-encoded reporter protein expression (violet-excited [VEX] or green fluorescent protein [GFP]), mixed at a 1:1 ratio and co-transferred into congenically distinct LCMV clone 13 (Cl13) infected mice (**Fig. 2A**). On d8p.i., expression of Tcf1 (or Ly108) and Tim3 can identify T_EX_ precursors (T_EX_^prec^; Tcf1/Ly108^+^Tim3^−^) or more differentiated “effector-like” CD8 T cells (Tcf1/Ly108^-^ Tim3^+^) (**Fig. S2A**).^33, 35^ Both d8 populations were detected for P14 Empty controls. In contrast, most of the P14 STAT5CA cells (91.7±0.98%) had differentiated into Ly108^−^ Tim-3^+^ CD8 T cells whereas the Ly108^+^Tim-3^−^ T_EX_^prec^ population was substantially reduced in both frequency (2.7±0.4%) and absolute number for the STAT5CA RV group (4.3-fold lower numbers than the Empty RV group) (**Fig. 2B** and **S2B**). We further confirmed the decrease in the generation of T_EX_^prec^ cells in P14 STAT5CA cells using unbiased clustering based on 12 flow cytometry parameters (**Fig. 2C**-see methods) or Tcf1 expression (**Fig. 2D**-upper panel). The overall numerical advantage of P14 STAT5CA cells over the P14 Empty at this early time-point (**Fig. S2B**) suggested a differentiation bias towards the Ly108^−^Tim-3^+^ effector-like population at the expense of T_EX_^prec^. Enforcing Stat5a activity also resulted in substantially lower Tox expression (**Fig. 2D-**lower panel). At this early time point, production of antiviral cytokines by P14 Empty and P14 STAT5CA cells after gp33 peptide stimulation was similar (**Fig. S2C**). Expression of the inhibitory receptors (IR) PD-1, Lag-3, and Tigit was also equivalent between STAT5CA and Empty RV P14 cells whereas other IRs (2B4, Tim-3) and some effector-related molecules (Cx3cr1, Granzyme B [GzmB]) were more highly expressed by the STAT5CA P14 cells (**Fig. 2E**). Thus, during the first week of a chronic viral infection, increasing STAT5a activity reduced Tox expression and enhanced the development of Ly108^−^Tim-3^+^ effector-like cells at the expense of T_EX_^prec^.

**Figure 2:**
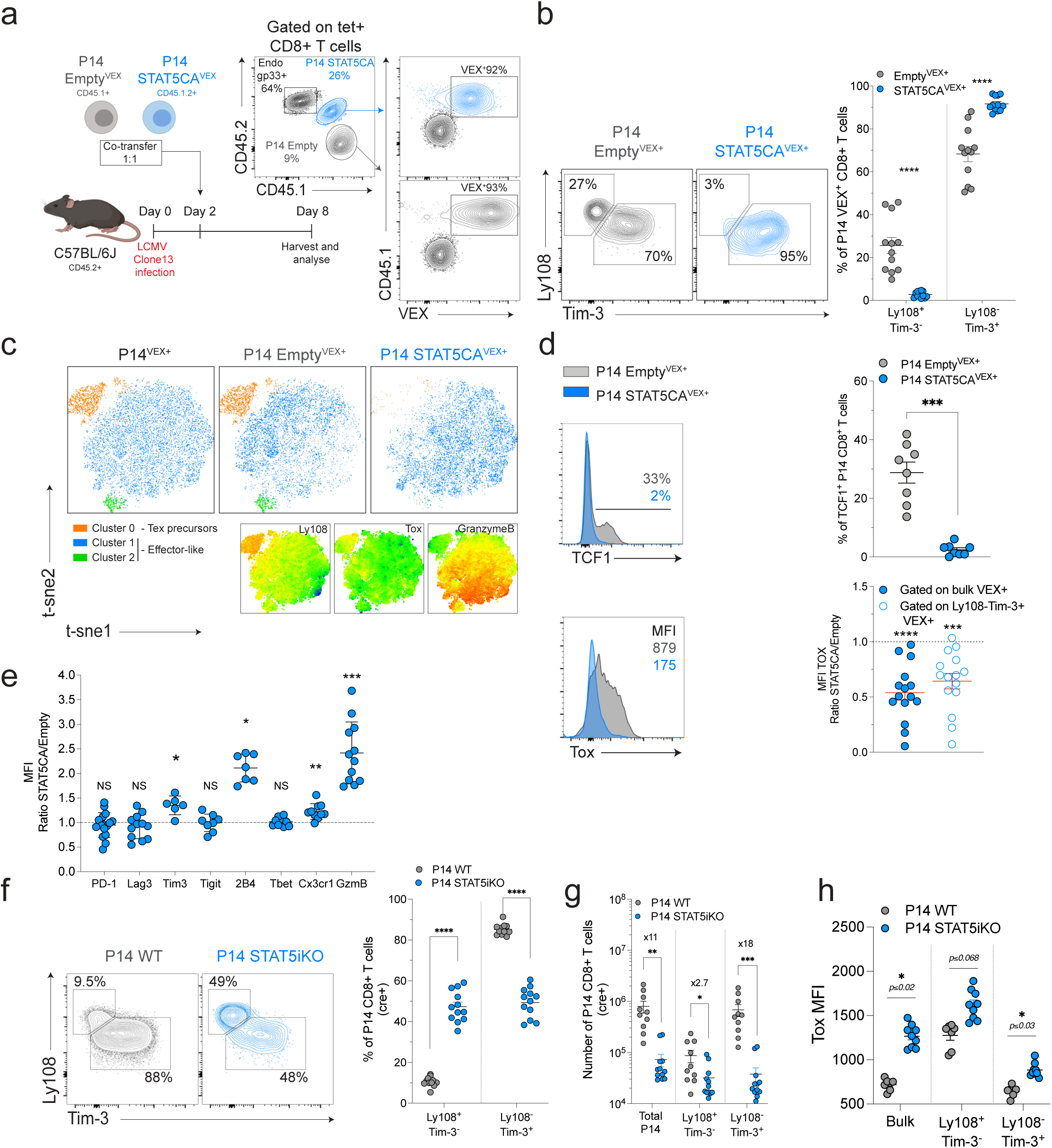
Stat5 opposes Tox and antagonizes establishment of exhaustion. **A**-Experimental design. **B**-Frequency of Ly108/Tim-3-defined subsets in indicated populations at d8p.i. **C**-t-sne representation of flow cytometry data highlighting FlowSOM clusters (see methods). **D**-Tcf1 and Tox expression in indicated populations at d8p.i. **E**-MFI of indicated markers expressed as a ratio (P14 STAT5CA/P14 Empty). **F**-Frequency of Ly108/Tim-3-defined subsets in indicated populations at d8p.i. **G**-Absolute numbers of indicated populations at d8p.i. **H**-Tox expression in indicated sub-populations of P14 WT and P14 Stat5iKO at d8p.i.

Stat5 supports early expansion of Ag-specific CD8 T cells in acutely resolving infections, though Stat5 activity in these settings has minimal impact on the balance of KLRG1^+^ terminal T_EFF_ and CD127^+^ memory precursors CD8 T cells.^38, 39^ To interrogate how Stat5 regulates differentiation of Ag-specific CD8 T cells early during chronic viral infection, naïve P14 WT (Rosa^YFP^*Stat5a/b*^+/+^) and P14 Stat5iKO (Rosa^YFP^*Stat5a/b*^fl/fl^) were treated with tat-cre recombinase *in vitro* (cre^+^; or not [cre^-^]) to induce genetic deletion of *floxed* alleles (**Fig. S2D**). Induction of YFP served as a surrogate for efficient cre-mediated recombination. We then adoptively transferred cre treated (cre^+^) P14 WT or P14 Stat5iKO cells (or their cre^-^ controls) into congenically distinct recipient mice that were infected with LCMV Cl13 and assessed the impact of loss of *Stat5a/b* at d8p.i. Genetic deletion of *Stat5a/b* resulted in a higher proportion of T_EX_^prec^ cells and a reduction in Ly108^−^Tim-3^+^ effector-like cells (11±1% vs 47±2% and 85±1% vs 49±2% of T_EX_^prec^ and Ly108^−^Tim-3^+^ cells in P14 WT vs P14 Stat5iKO respectively) (**Fig. 2F** and **S2E-F**). The overall number of P14 cells was reduced by ∼11-fold in the absence of *Stat5a/b* but this effect was predominantly in the Ly108^−^Tim-3^+^ population that was reduced ∼18-fold compared to the WT P14 cells (**Fig. 2G**). In contrast, the Ly108^+^Tim-3^−^ T_EX_^prec^ were impacted less with only a ∼2.7-fold reduction in the absence of *Stat5a/b*. Changes in proliferation or cell death did not appear to explain these differences between the WT and *Stat5a/b* iKO cells because BrdU incorporation (d7-8) and active caspase 3 were similar between these populations (**Fig. S2G-H**). These data suggested an impact on differentiation and in particular, an altered formation of Ly108^−^Tim-3^+^ cells in the absence of *Stat5a/b*, consistent with the enhanced development of this effector-like population using STAT5CA (**Fig. 2B**). This effect of *Stat5a/b*-deficiency appeared to be preferential to chronic infection because, consistent with previous studies,^38, 39^ there was little impact of loss of *Stat5a/b* on the distribution of memory precursor and short-lived effector CD8 T cell subsets during acute LCMV Armstrong (Arm) infection (**Fig. S2I**). Consistent with the reduction of Tox upon constitutive activation of Stat5a (**Fig. 2D**), loss of *Stat5a/b* resulted in increased Tox expression in P14 Stat5iKO compared to P14 WT during Cl13 infection (**Fig. 2H**). The minimal role of Tox at early stages of acutely resolving versus chronic infection^6,7^ may explain the preferential impact of Stat5 during LCMV Cl13 infection compared to Arm infection. Moreover, these data indicate a role for Stat5 in the early population dynamics during evolving chronic infection with Stat5 activity repressing Tox and fostering differentiation into the more effector-like Ly108^−^Tim-3^+^ subset during the first week of infection.

### Constitutive STAT5a activation promotes an epigenetic state with features of effector and exhausted CD8 T cells during chronic infection

Commitment of CD8 T cells to exhaustion is associated with acquisition of a distinct epigenetic landscape.^6,26, 40^ To investigate whether constitutively active Stat5a altered early epigenetic programming of Ag-specific CD8 T cells, we performed <u>A</u>ssay for <u>T</u>ransposase-<u>A</u>ccessible <u>C</u>hromatin followed by high throughput sequencing (ATACseq)^41^ on P14 Empty and P14 STAT5CA cells at d8 of Cl13 infection and compared these data to P14 cells isolated at d8 of Arm infection (T_EFF_). Examining all differentially accessible peaks (DAPs; lfc>2, FDR≤0.01), each population of P14 cells was distinct in principal component space (**Fig. 3A**) or by Spearman distance analysis (**Fig. S3A**) with P14 Empty and P14 STAT5CA differing from each other by almost the same “distance” as they each differed from the T_EFF_ cells from Arm infection. Indeed, P14 Empty and P14 STAT5CA differed from Arm-derived T_EFF_ by 27,876 and 25,681 DAPs respectively. P14 STAT5CA cells also differed from their P14 Empty controls in chronic infection by 16,901 DAPs (**Fig. 3B, Table S2**). The majority of DAP were at intergenic, promoter and intronic regions (**Fig. S3B**). One possible explanation for these epigenetic differences could be the near absence of the Ly108^+^Tcf1^+^ T_EX_^prec^ population in P14 STAT5CA (**Fig. 2B**) and subsequent absence of the Tcf1 signature. However, of the 16,901 DAPs between P14 Empty and P14 STAT5CA cells, only 454 peaks (2.7%) related to genes differentially expressed between WT and *Tcf7*^-/-^ gp33^+^ CD8 T cells from d7 post LCMV Cl13 infection (**Fig. S3C)**.^33^ This observation suggested that the distinct ATAC profile of P14 STAT5CA cells was not simply due to the absence of Tcf1-expressing T_EX_^prec^ cells in this population, but rather reflected epigenetic remodeling provoked by constitutive Stat5a expression.

**Figure 3:**
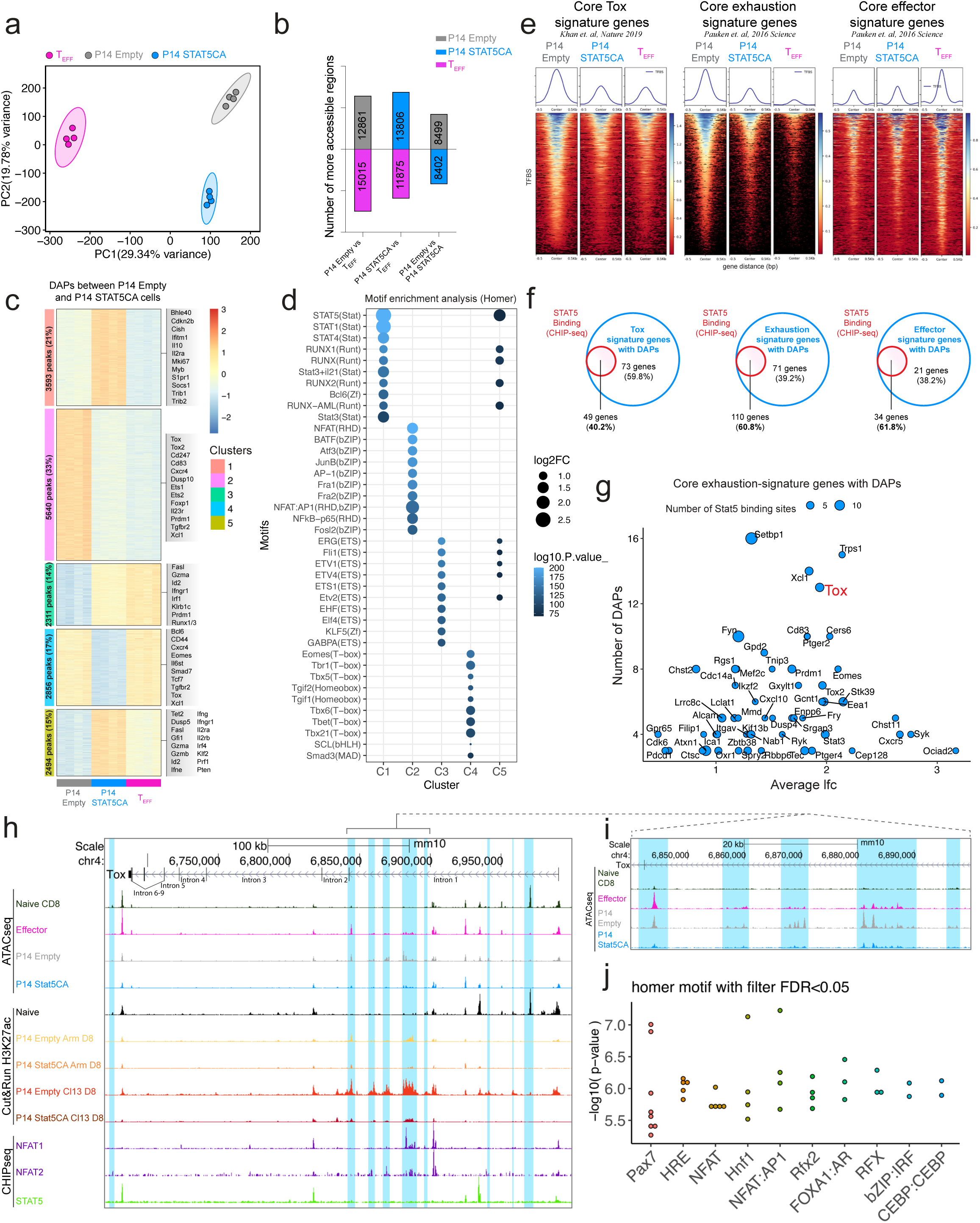
Enhanced Stat5a activity restrains Tox and the Tox-dependent exhaustion program while supporting an effector epigenetic landscape. **Splenic P14 Empty and** P14 STAT5CA cells isolated at d8 of Cl13 infection were analyzed by ATACseq. Naïve CD8 T cells (not depicted) and P14 cells isolated from Arm infected mice at d8p.i. (T_EFF_) were used as reference. **A**-PCA of normalized ATACseq counts (all peaks). **B**-Number of peaks more accessible in indicated populations and comparisons (FDR 0.01, lfc≥2). **C**-Clustered heatmap (k-means) of DAPs between P14 Empty and P14 STAT5CA (FDR 0.01, lfc≥2) annotated with most variable genes per cluster (among top 100 per cluster by lfc and number of DAPs). **D**-Top 10 motifs (Homer) enriched in DAPs from corresponding clusters in Fig.3C. **E**-Heatmap for scores associated with genomic regions plotting accessibility at genes from indicated gene signatures list. **F**-Frequency of genes from signature list in Fig. 3E with DAPs between P14 Empty and P14 STAT5CA that possess direct binding sites for Stat5 (chipseq dataset from *Villarino et. al, J Exp Med 2017*).^84^ **G**-Dot plot of exhaustion signature genes (**from** Fig. 3E-middle) containing at least 3 DAPs between P14 Empty and P14 STAT5CA cells and scored based on the number of DAPs per gene (Y axis), average lfc (X axis) and number of direct Stat5 binding sites (bubble size). **H**-ATACseq, Cut&Run (H3K27ac) and Chip-seq (NFAT1,^42^ NFAT2^43^ and Stat5^84^) tracks at the Tox locus. Blue highlights indicate ATAC peaks reduced in P14 STAT5CA cells compared to P14 Empty. **I**-ATACseq track zoom-in from Fig. 3H. **J**-Top 10 motifs (Homer) enriched in DAPs between P14 Empty and P14 STAT5CA found at the Tox locus.

To examine how constitutive Stat5 expression impacted the early establishment of exhaustion in Ag-specific CD8 T cells, we performed K-means clustering of the 16,901 DAPs between P14 Empty and P14 STAT5CA (**Fig. 3C, Table S2**). Cluster 1 and 4 (C1,- 4) contained regions preferentially remodeled in P14 STAT5CA cells. In contrast, C2, -3 and -5 contained DAPs with selectively increased or decreased accessibility in P14 Empty cells but the opposite trend in both P14 STAT5CA and T_EFF_ cells (**Fig. 3C**). Hence, among the 16,901 DAPs between P14 Empty and P14 STAT5CA cells, 62% (10,445 peaks; [C2,-3 and-5]) represented changes in which P14 STAT5CA cells became more similar to T_EFF_ compared to P14 Empty cells. DAPs in C1 (gained in P14 STAT5CA) strongly enriched for Stat5 binding motifs (also potentially bound by Stat1, -3 and -4) and motifs for effector-related TFs (Runx1/2) (**Fig. 3C-D**; **Table S2**). C4 DAPs (decreased in P14 STAT5CA) rather contained T-box and homeobox TFs motifs (i.e. Eomes, T-bet, Tbr1, Tbx2-6, Tgif1/2). C2 with high accessibility in P14 Empty was enriched in binding motifs for TCR-dependent TFs with established roles in T_EX_ including NFAT (RHD) and BATF (bZIP) and this cluster included open chromatin regions at exhaustion-specific genes (i.e. *Tox*, *Tox2*) (**Fig. 3C-D** and **S3D**). C3 and -5 with increased accessibility in both P14 STAT5CA and T_EFF_ compared to P14 Empty, contained several DAPs located near genes encoding effector-related molecules and TFs (i.e. *Gzma*, *Fasl*, *Prf1*, *Runx1/3*, *Id2*) and enriched for Stat5, Runx1/2 and ETS motifs (**Fig. 3C-D** and **S3D**). Further clustering of all DAPs between P14 Empty, P14 STAT5CA and T_EFF_ (70,458 peaks) revealed decreased accessibility at a large fraction of exhaustion-related open chromatin regions in P14 STAT5CA cells concomitant with a shift to a more effector-like open chromatin landscape (**Fig. S3E**). Together, these data highlight an altered epigenetic program in P14 STAT5CA cells early during chronic viral infection and shift towards effector biology at the stage of differentiation when epigenetic imprinting of exhaustion typically occurs.

Consistent with this antagonism of the early program of exhaustion, a large fraction of open chromatin regions near Tox-dependent or exhaustion-related genes were lost in P14 STAT5CA cells, whereas accessibility at effector-associated genes was increased (**Fig. 3E**). A substantial proportion of genes from these core signature lists possessed one or several Stat5 binding sites, suggesting direct regulation of genes involved in effector versus exhaustion biology by this TF (**Fig. 3F**, **Table S2**). *Tox* was among the top exhaustion-related genes with reduced chromatin accessibility in P14 STAT5CA compared to P14 Empty cells consistent with lower expression of Tox in the former population (**Fig. 2D**). Moreover, this gene contained one of the highest numbers of direct Stat5 binding sites (**Fig. 3G**). To explore the relationship between Tox and Stat5a, we further analyzed the *Tox* locus (**Fig. 3H**). Overall, chromatin accessibility decreased in *Tox* in P14 STAT5CA compared to P14 Empty cells, notably at enhancers in the first intron (**Fig. 3H-I**). Indeed, accessibility at one region in the first intron was decreased to a level comparable to or even lower than that observed in T_EFF_ cells (**Fig. 3I**). This region was also enriched for active H3K27Ac histone marks, particularly in settings of chronic Ag-stimulation (P14 Empty Cl13 D8) suggesting an active chromatin environment in early T_EX_ (**Fig 3H**). Indeed, this region contained, and was framed by several binding sites for NFAT1^42^ and NFAT2^43^ (**Fig. 3H,J**), TFs that are key drivers of Tox induction during the early development of T ^6, 7, 10^ This active H3K27Ac chromatin environment in intron 1 was reduced in P14 STAT5CA cells suggesting that enforced STAT5 activity impedes chromatin accessibility in *Tox*, particularly at sites where the transcriptional drivers of Tox induction (NFATs) can bind. Together, these data highlight the potential impact of Stat5 in the epigenetic regulation of key exhaustion-related genes including *Tox* where Stat5 appears to function directly to modulate accessibility in this locus at locations where NFAT proteins may act.

### Constitutive STAT5a activation instigates a distinct effector/NK-like transcriptional program and improves therapeutic potential

Establishment of a Tox-dependent exhaustion program drives altered function, but also is required for the maintenance of T_EX_ during chronic infections and cancer.^6–9, 31, 32^ Without Tox, T_EX_ cannot form or persist in the setting of chronic Ag stimulation. Because constitutive Stat5a activation antagonized Tox, we next investigated the durability and fate of P14 STAT5CA cells later during chronic viral infection. We again employed the co-adoptive transfer model where P14 Empty and P14 STAT5CA cells could be examined in the same chronically infected recipient mice. At d27p.i., 85±4.5% of the RV transduced (VEX^+^) donor P14 cells were P14 STAT5CA cells and this population numerically outcompeted their P14 Empty counterpart in the spleen as well as peripheral tissues (**Fig. 4A-B** and **S4A-C**). This numerical advantage of P14 STAT5CA cells was also observed even in mice with life-long viremia due to transient depletion of CD4 T cells (Cl13 αCD4; **Fig. 4A**-right) where cells persisted for at least ∼3 months (**Fig. S4D**) suggesting a prolonged advantage of this enforced STAT5CA expression. At d27p.i., the donor P14 STAT5CA population was enriched for Ly108^−^CD69^−^ T_EX_^int^ cells. The proportion of the two Tcf1^+^ progenitor subsets, T_EX_^prog1^ (Ly108^+^CD69^+^) and T_EX_^prog2^ (Ly108^+^CD69^−^) were dramatically reduced, though a small population of the latter was present, and the frequency of T_EX_^term^ (Ly108^−^CD69^+^) was also reduced compared to the P14 Empty population from the same mice (**Fig. 4C** and **S4E**). Thus, constitutive Stat5a activation leads to an accumulation advantage for virus-specific CD8 T cells in the setting of a chronic viral infection, despite the substantial reduction in Tcf1^+^ T_EX_^prog^ cells, mainly through an accumulation of T_EX_^int^-like cells.

**Figure 4:**
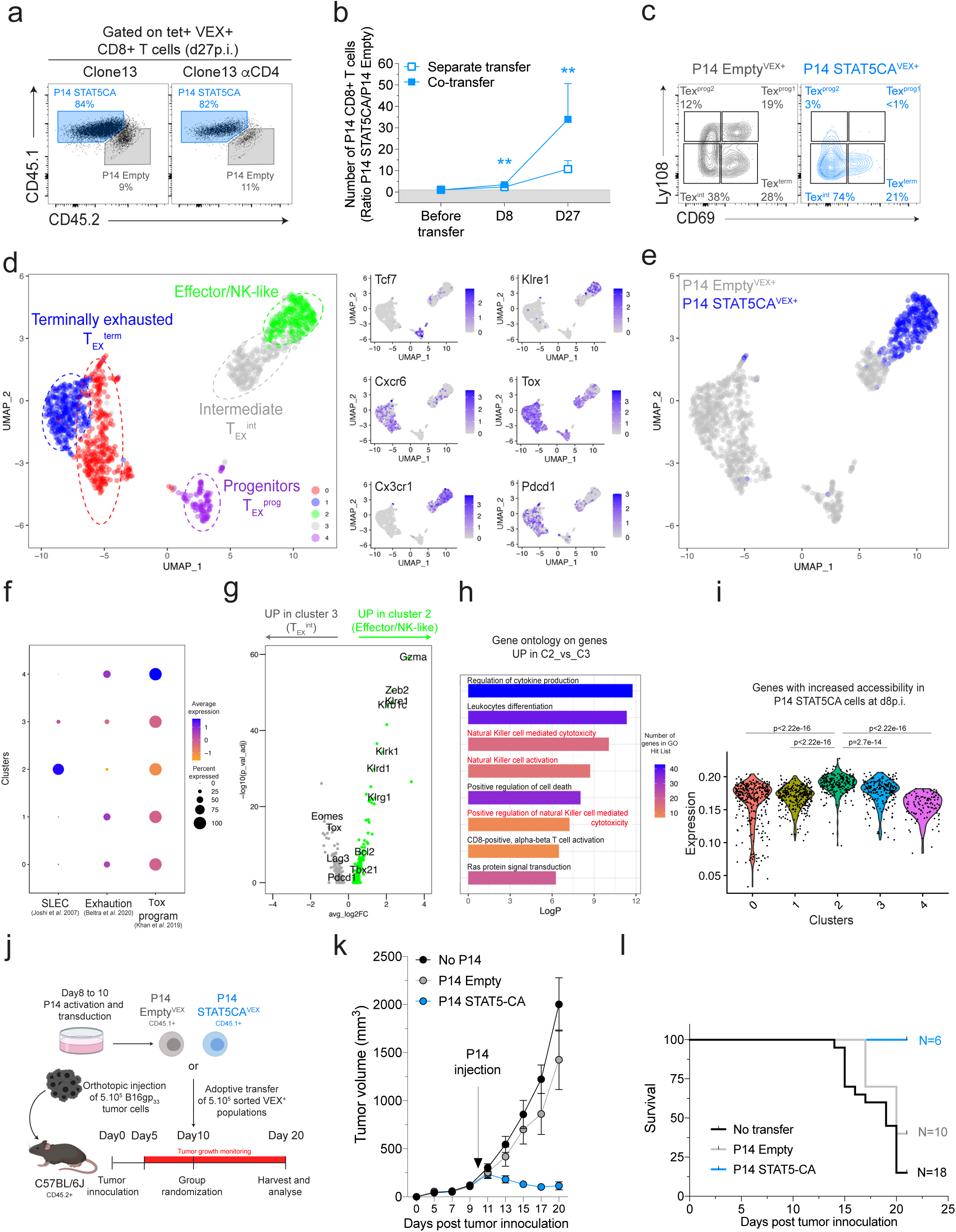
Constitutive Stat5a activity drives a durable and protective effector/NK-like CD8 T cell differentiation during chronic viral infection and cancer. **A-** Frequency of co-transferred P14 Empty and P14 STAT5CA cells among RV+ (VEX+) CD8 T cells at d27p.i. with Cl13 (left) or Cl13 with CD4-depletion (right). B-Ratio of cell number (P14 STAT5CA/P14 Empty) at indicated time points in the spleen. C-Frequency of Ly108/CD69-defined T_EX_ subsets in indicated populations at d27p.i. D-E UMAP of scRNAseq data combining P14 Empty^VEX+^ and P14 STAT5CA^VEX+^ cells isolated at d27p.i. plotting Seurat clusters (D-left) or individual samples (E) and representative genes per cluster (D-right). F-GSEA for indicated signatures in Seurat clusters. G-DEGs (log2FC>0.5, p_value_adj≤0.05) between C2 and C3 from Fig. 4D. H-Gene ontology for genes Up in C2 vs C3 from (Fig. 4D) I-Relative expression of genes with increased accessibility in P14 STAT5CA over P14 Empty at d8p.i. (Fig. 3) across Seurat clusters. J-Experimental design. K-L-B16_gp33_ tumor growth (K) and Kaplan Meyer survival curve (**L**) for each experimental group.

Based on the ATAC-seq data from d8 p.i., we hypothesized that the P14 STAT5CA cells later in chronic infection might differ from previously defined T_EX_^int^ despite their Ly108^−^ CD69^−^ phenotype. We therefore performed single-cell RNA sequencing (scRNA-seq) to compare P14 STAT5CA and P14 Empty cells at ∼1 month of chronic viral infection. We identified canonical clusters of T_EX_ cells including a T_EX_ progenitor cluster (C4; progenitors [T_EX_^prog^]) selectively expressing *Tcf7, Slamf6,* and *Xcl1* and two clusters that enriched for a signature of T_EX_^term^ (C0 and C1) and expressed *Cxcr6*, a marker associated with terminal exhaustion^30^ as well as elevated *Pdcd1*, *Cd160 and CD244* (2B4) (**Fig. 4D** and **S4F-G**). We also identified two clusters of *Cx3cr1*-expressing cells, one of which was consistent with T_EX_^int^ cells (C3). A second cluster (C2; Effector/NK-like) also enriched for a T_EX_^int^ cell signature but displayed selective expression of NK cell receptors (i.e. *Klre1*, *Klrk1*, *Klrd1*), elevated transcripts of effector molecules and TFs (i.e. *Gzma, Zeb2*) (**Fig. 4D** and **S4F-G, Table S3**), similar to a recently described NK-like T_EX_ subset.^26, 44^ This subset also had reduced expression of several exhaustion-related genes including *Tox* and *Pdcd1* compared to all other clusters (**Fig. 4D** and **S4F-G, Table S3**). This effector/NK-like cluster (C2) was composed mostly of P14 STAT5CA cells with little contribution of P14 Empty cells. Rather, the latter were more evenly distributed throughout C0,1,3, and -4 (**Fig. 4E** and **S4H**). Consistent with the high expression of NK- and effector-genes in C2, these cells also enriched for the transcriptional signatures of short-lived effector CD8 T cells (SLEC) and had the lowest enrichment score for an exhaustion or Tox-dependent transcriptional signature among clusters of CD8 T cells from Cl13 infection (**Fig. 4F**). Moreover, although the P14 STAT5CA cluster (C2) mapped closely to the T_EX_^int^ cluster (C3) in a UMAP representation (**Fig. 4D**), these two clusters differed by expression of 211 genes (Log2FC>0.5, p_value_adj≤0.05) with T_EX_^int^ cells mostly composed of P14 Empty cells having higher expression of exhaustion-related genes including *Tox*, *Eomes*, *Pdcd1, Lag3* whereas the P14 STAT5CA cells had higher expression of genes encoding effector or NK cell-related markers (e.g. *Klre1, Klrb1c, Klrk1, Klrd1*, *Gzma, Tbx21*), and genes involved in cell survival (e.g. *Bcl2)* (**Fig. 4G-H, Table S3**). Lastly, genes with increased chromatin accessibility in P14 STAT5CA cells at d8p.i. (**Fig. 3, Table S4**) had higher mRNA expression in P14 STAT5CA cells (C2) at d27p.i. (**Fig. 4I**), suggesting that the transcriptional differences observed in P14 STAT5CA cells at d27p.i. reflected, at least in part, the chromatin accessibility landscape established by d8 p.i. Together these data suggested that constitutive STAT5a activation drove virus-specific CD8 T cells into a distinct state during chronic infection characterized by a transcriptional program with both effector and NK-like features in this setting that typically drives CD8 T cell exhaustion.

The enhanced effector/NK biology of P14 STAT5CA cells coupled with the accumulation advantage in chronic infection prompted us to evaluate disease control and/or therapeutic efficacy. Because during LCMV Cl13 infection even small changes in the number of WT P14 alter pathogenesis^45, 46^ complicating questions of protective immunity, we used a tumor model where the ability to control tumor growth could be assessed. P14 STAT5CA and P14 Empty cells were adoptively transferred separately into mice with established B16-gp_33-41_ tumors (d10 post tumor inoculation) (**Fig. 4J**). Adoptive transfer of P14 Empty cells into these mice only slightly delayed tumor growth whereas P14 STAT5CA cells resulted in substantial reduction in tumor burden and survival of all mice in this group (**Fig. 4K-L**). Thus, the impact of constitutively active STAT5a on differentiation of CD8 T cells not only antagonized exhaustion, but these changes in CD8 T cell differentiation corresponded to improved therapeutic efficacy and control of tumor growth.

### STAT5 is essential for generation of T_EX_^int^ cells and for response to PD-L1 blockade

We next used the adoptive transfer approach to investigate how endogenous Stat5 influenced T_EX_ dynamics when exhaustion was fully established using P14 WT and P14 Stat5iKO cells (**Fig. S2D**). At ∼1 month p.i., P14 Stat5iKO cells had high expression PD-1 and Tox (albeit slightly reduced compared to P14 WT) but lacked expression of molecules associated with T_EX_^int^ or T_EX_^term^ (i.e. Granzyme B, Cx3cr1, Tim-3; **Fig. 5A**). Indeed, the P14 Stat5iKO population was almost exclusively composed of T_EX_ progenitors (T_EX_^prog1^ and T_EX_^prog2^) with only minor populations of T_EX_^int^ cells and T_EX_^term^ compared to P14 WT cells or STAT5-proficient (cre-/YFP-) controls from the same donor P14 population (**Fig. 5B** and **Fig. S5A-B**). As a result, the number of P14 Stat5iKO cells in the spleen was reduced compared to P14 WT cells with even more substantial reductions in the blood and peripheral tissues consistent with the accumulation of T_EX_^int^ and T_EX_^term^ cells in these locations (**Fig. 5C** and **S5C-D**).^27^ However, the T_EX_^prog1^ and T_EX_^prog2^ compartments remained numerically intact in the absence of *Stat5a/b* (**Fig. 5C**). The stability of these progenitor-like CD8 T cells in chronic viral infection in the absence of *Stat5a/b* was in stark contrast to the ∼23-fold reduction in the number of memory CD8 T cells formed by P14 Stat5iKO following an acute infection with LCMV Arm (**Fig. 5D**) highlighting the distinct dependency on *Stat5a/b* for the formation and/or maintenance of T_MEM_ in acutely resolving versus T_EX_^prog^ in chronic viral infection.

**Figure 5:**
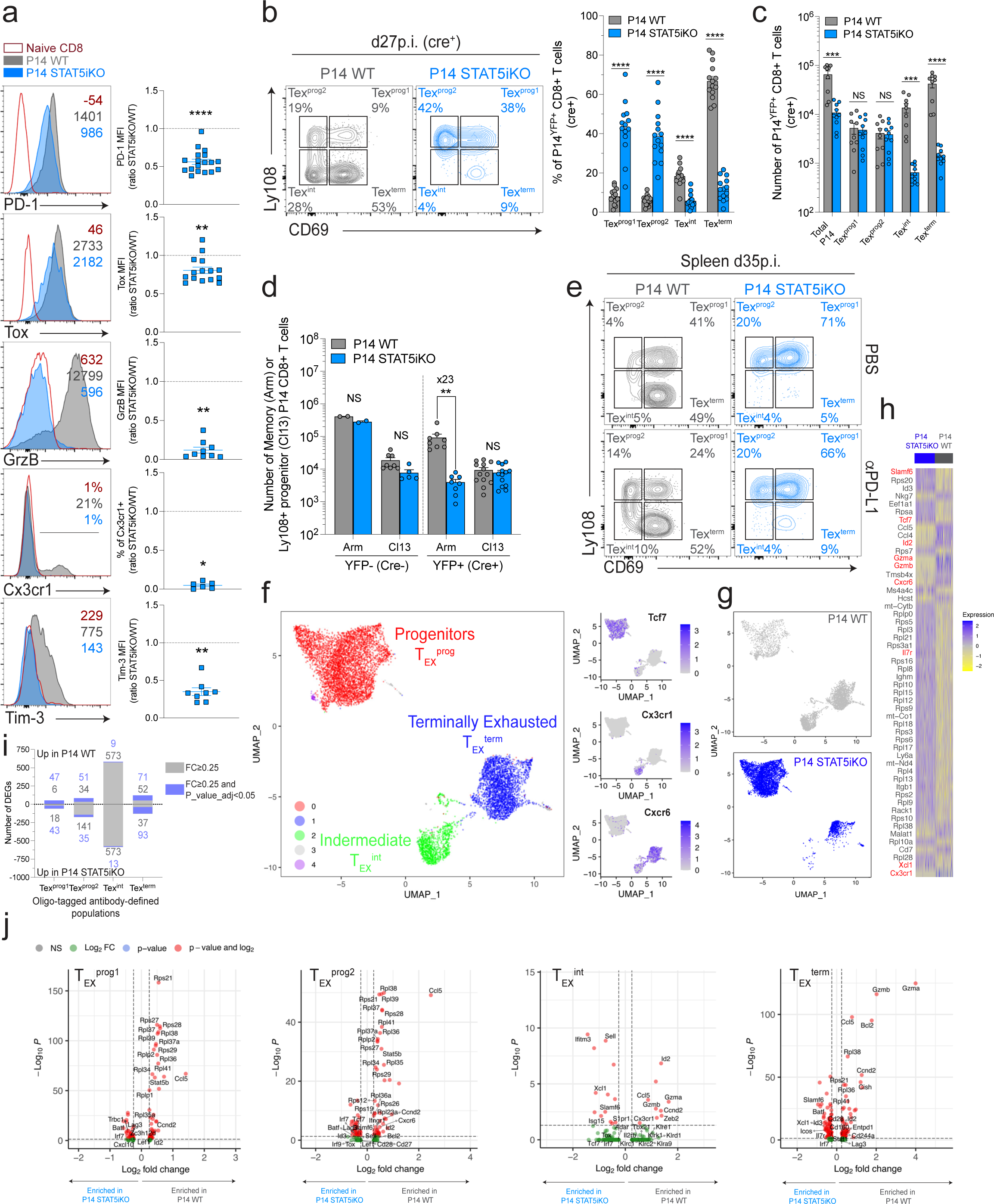
Stat5-signals drive T_EX_^int^ cell development and are essential for CD8 T cell responses to PD-L1 blockade. **A-** Expression of key markers on indicated splenic populations at d27p.i. **B**-Frequency of Ly108/CD69-defined subsets among indicated populations at d27p.i. **C-** Absolute numbers of indicated populations of P14 WT and P14 Stat5iKO cells at d27p.i. **D**-Absolute numbers of T_MEM_ and Ly108^+^ T_EX_^prog^ in indicated P14 populations at d27 post Arm (Arm, memory) or Cl13 (Cl13, Ly108^+^ progenitors) infection with (YFP^+^cre^+^) or without (YFP^-^cre^-^) prior *in vitro* treatment with tat-cre recombinase. **E**-Frequency of Ly108/CD69-defined subsets among P14 WT and P14 Stat5iKO cells at d35p.i. in CD4 T cells-depleted hosts treated (αPD-L1) or not (PBS) with anti-PD-L1 antibodies between d22-34 (see method). **F**-**G**-UMAP plotting RNA-defined Seurat clusters (**F**-left) or individual samples (**G**) from CITE-seq analysis of P14 WT and P14 Stat5iKO cells isolated at d27p.i. **H**-Top 52 DEGs between P14 WT and P14 Stat5iKO. **I**-Number of DEGs between oligo-tagged antibodies (Ly108 and CD69)-defined populations (see **Fig. S6D,E**). **J**-DEGs (FC≥0.25) between indicated oligo-tagged antibodies (Ly108 and CD69)-defined populations of P14WT and P14 Stat5iKO cells.

As T_EX_^prog1^ cells exit quiescence, they convert to T_EX_^prog2^ cells that re-engage cell-cycle and further differentiate into T_EX_^int^ cells. This T_EX_^prog2^ to T_EX_^int^ transition is amplified by PD-1 pathway blockade.^27^ In the absence of *Stat5a/b*, proliferation of T_EX_^prog1^ and T_EX_^prog2^ was reduced compared to P14 WT (**Fig. S5E**). This reduced proliferation suggested a defect in the conversion of T_EX_^prog1^ and T_EX_^prog2^ into T_EX_^int^ cells without *Stat5a/b*. PD-L1 blockade did not rescue the development of T_EX_^int^ cells for P14 Stat5iKO cells and these cells failed to expand in number following PD-L1 blockade despite a numerically intact progenitor compartment (**Fig. 5E** and **S5F-G**). Thus, Stat5 was essential for proliferation-driven conversion of T_EX_^prog1^ and T_EX_^prog2^ into T_EX_^int^, a key developmental transition for generation of more terminally differentiated T_EX_ subsets, replenishment of peripheral immunity and response to PD-1 blockade.

To investigate the molecular effects of Stat5-deficiency in T_EX_ subsets, we performed Cellular Indexing of Transcriptomes and Epitopes by sequencing (CITE-seq) on P14 WT and P14 Stat5iKO cells at ∼1 month of chronic infection. Using RNA-based unsupervised clustering, we again identified major clusters of T_EX_ cells and confirmed the near absence and robust reduction of the T_EX_^int^ and T_EX_^term^ subsets respectively in P14 Stat5iKO compared to P14 WT (**Fig. 5F,G** and **S6A,B**). Top DEGs between WT and Stat5iKO P14 cells reflected this altered subset distribution with increased expression of progenitor-associated genes (i.e. *Tcf7*, *Id3*, *Xcl1*) but depletion of genes related to more differentiated subsets (i.e. *Cx3cr1*, *Cxcr6*, *Gzma*, *Gzmb*) in the later population (**Fig. 5H, Table S5**). Comparing clusters also revealed transcriptional divergence between P14 Stat5iKO and P14 WT cells in the three main clusters, suggesting that STAT5-deficiency affected all major subsets of T_EX_ cells (**Fig. S6C**).

The T_EX_^prog1^ and T_EX_^prog2^ often co-segregate in scRNA-seq space because of the dominance of the progenitor signature. However, the T_EX_^prog2^ subset engages distinct biology as these cells begin to downregulate *Tcf7*, enter cell cycle and initiate the transition to downstream T_EX_ subsets.^27^ CITE-seq captured this key transitional biology by discriminating Ly108^+^CD69^−^ T_EX_^prog2^ cells using surface markers and allowed us to further interrogate Stat5a/b-dependent transcriptional differences in this subset (**Fig. S6D-F**). Indeed, analysis of DEGs confirmed transcriptional differences between oligo-tagged antibody-defined T_EX_ subsets (**Fig. 5I, Table S5**). Of interest, the Stat5-dependent cyclin *Ccnd2* that initiates the G1-S cell-cycle phase was elevated in all P14 WT T_EX_ subset compared to P14 Stat5iKO cells consistent with a defect in cell-cycle re-entry in the absence of *Stat5a/b* (**Fig. 5J**). This reduced *Ccnd2* expression was coupled with a robust decrease in expression of multiple genes encoding ribosomal proteins (*Rps* genes) in P14 Stat5iKO cells, particularly in T_EX_^prog1^ and T_EX_^prog2^ cells (**Fig. 5J**). This observation suggested a reduction in protein synthesis in *Stat5a/b*-deficient T_EX_ coupled to a defect in differentiation potential. *Stat5a/b*-deficient T_EX_^prog2^ cells also retained higher expression of progenitor associated-molecules (i.e. *Tcf7*, *Sell*, *Slamf6*, *Id3*, *Tox*) whereas these genes are typically reduced during the T_EX_^prog1^ to T_EX_^prog2^ transition.^27^ Expression of some of these progenitor-related molecules even trended higher in the few *Stat5a/b*-deficient T_EX_^int^ and T_EX_^term^ cells detectable (e.g. *Sell*, *Slamf6*, *Tcf7*, *Tox*) compared to P14 WT T_EX_^int^ and T_EX_^term^ cells (**Fig. 5J**). In addition, the few T_EX_^int^ cells that developed in the absence of *Stat5a/b* had impaired expression key effector genes including (i.e. *Gzma*, *GzmB*, *Cx3cr1*, *Zeb2*, *Tbx21*, *Id2*) and multiple KLR-molecules (i.e. *Klrd1*, *Klre1*, *Klrc1*, *Klrc2*, *Klrk1*), though because of the small number of Stat5iKO T_EX_^int^ cells, many of these changes did not reach statistical significance. However, this lack of effector-biology in P14 Stat5iKO T_EX_^int^ cells was also apparent at the T_EX_^term^ stage (i.e. reduced *Gzmb*, *Gzma*) and these cells also had reduced expression of *Bcl2* perhaps contributing to poor survival in the absence of *Stat5a/b* (**Fig. 5J**). Together, these data point to a key inability of T_EX_^prog2^ cells to exit quiescence, re-engage cell-cycle and engage the transition to the T_EX_^int^ stage in the absence of *Stat5a/b*. Stat5 also mediated the transcriptional switch that accompanied this T_EX_^prog2^-to-T_EX_^int^ transition by extinguishing at least some of the T_EX_ progenitor-associated biology and fostering the effector and NK-like features that characterize the T_EX_^int^ subset.

### Temporal reactivation of Stat5 in T**_EX_** cells drives T**_EX_**^int^ cell accumulation and synergizes with PD-L1 blockade

Given the key role of Stat5 for T_EX_^int^ cell generation described above, we next explored the potential of temporally manipulating this axis to foster development of this subset. To this end, we leveraged an orthogonal IL-2/IL2Rβ system.^47^ Briefly, P14 CD8 T cells were transduced with an RV encoding an orthogonal IL2Rβ-receptor chain (*ortho*IL2Rβ; P14 IL2Rβ-ortho) that selectively binds and triggers Stat5 activation in response to cognate orthogonal IL-2 (*ortho*IL-2) but not the native endogenous IL-2 and compared these cells to those transduced with an empty RV (P14 Empty). Congenically distinct P14 IL2Rβ-ortho and P14 Empty cells were co-transferred into mice infected with LCMV Cl13 (**Fig. 6A**). Starting on d21p.i., groups of mice received escalated doses of *ortho*IL-2 for 5 days and changes in total T_EX_ and T_EX_ subsets were examined at d26p.i. *In vivo* delivery of *ortho*IL-2 caused a selective and dose dependent expansion of the YFP^+^ (RV^+^) P14 IL2Rβ*-*ortho cells compared to their P14 Empty control counterparts (**Fig. 6B**). This dose-dependent numerical increase was not observed in YFP^-^ (RV^-^) cells and, unlike WT IL-2, *ortho*IL-2 treatment also did not alter the frequency of regulatory T cells, demonstrating the specificity of the *ortho*IL-2/IL2Rβ system (**Fig S7A-B**). Moreover, although the frequency of T_EX_ subsets remained similar in the P14 Empty population and endogenous gp_33-41-_specific CD8 T cells (**Fig. S7C-D**), the P14 IL2Rβ-ortho population in the same mice displayed an expansion of T_EX_^prog2^ and T_EX_^int^ cells and concomitant reductions in the frequency of T_EX_^prog1^ and T_EX_^term^ cells with increasing doses of *ortho*IL-2 in both the spleen and peripheral tissues (**Fig. 6C-D** and **S7E-H**). The gradual decrease in T_EX_^term^ cell frequency within P14 IL2Rβ-ortho also suggested that *ortho*IL-2-mediated Stat5 activation stabilized the T_EX_^int^ stage, restraining conversion to the more terminally exhausted T_EX_^term^ cells. Although the frequency of stem-like T_EX_^prog1^ cells among the P14 IL2Rβ-ortho populations treated with *ortho*IL-2 decreased in a dose dependent manner, the absolute number of these key progenitor cells remained stable (**Fig. S7I**) suggesting that *ortho*IL-2 treatment could enhance the generation of downstream T_EX_ subsets without depleting the T_EX_ progenitor populations. These data indicate that temporal engagement of the *ortho*IL-2 system in T_EX_ cells, likely through increasing Stat5 activity, can function as an amplifier of the T_EX_^prog2^ transition into T_EX_^int^ cells.

**Figure 6:**
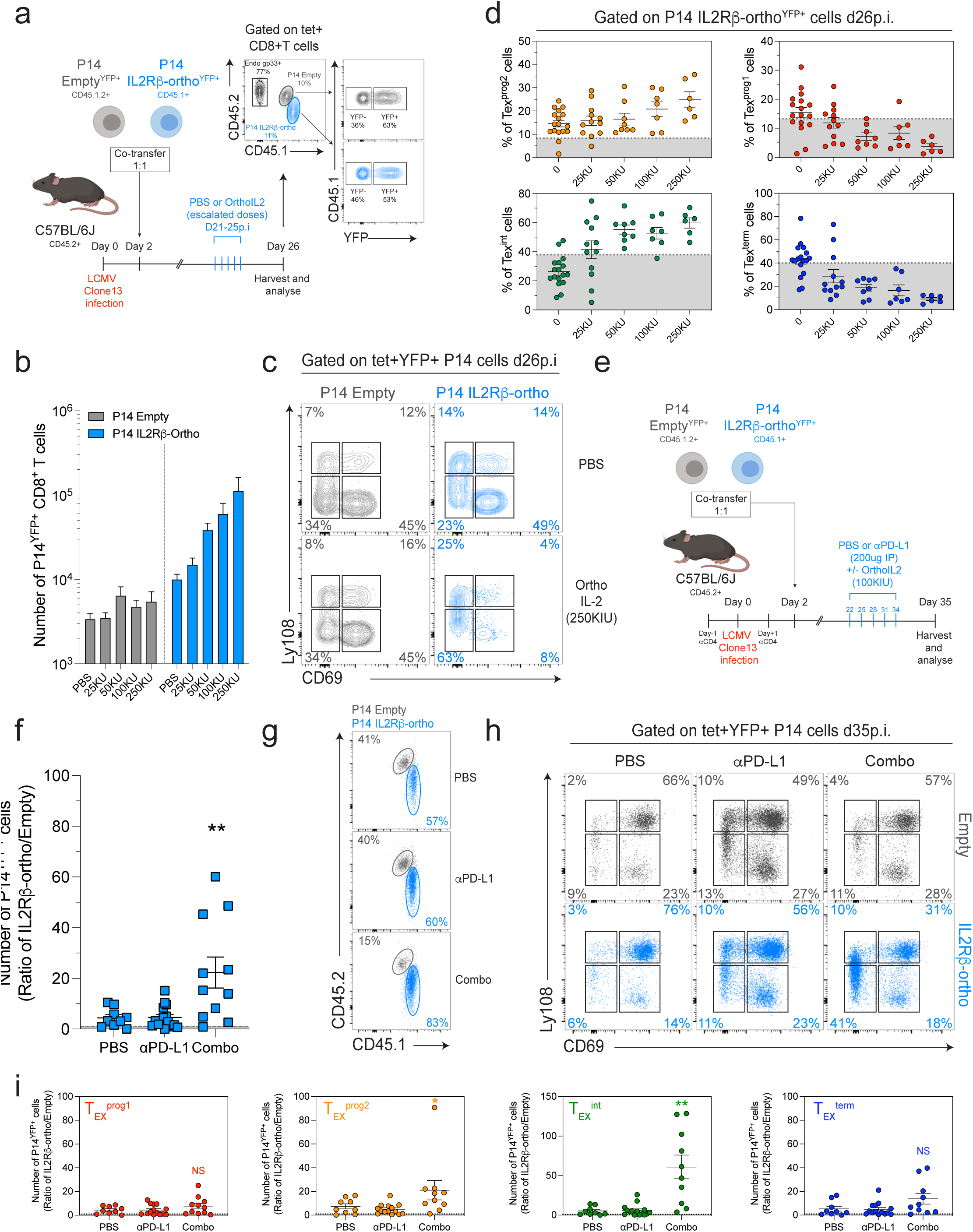
Orthogonal IL-2/IL2Rβ-triggered Stat5 activation in Ag-specific CD8 T cells enforces T_EX_^int^ cell development and synergizes with PD-L1 blockade. **A-** Experimental design. B-Absolute numbers of YFP^+^ P14 Empty and P14 IL2Rβ-ortho cells isolated at d26p.i. from experimental groups infused with indicated concentration of *ortho*IL-2. C-Frequency of Ly108/CD69-defined subsets among co-transferred P14 Empty^YFP+^ and P14 IL2Rβ-ortho^YFP+^ cells isolated at d26p.i. from indicated experimental groups. D-Frequency of indicated subsets among P14 IL2Rβ-ortho^YFP+^ cells isolated at d26p.i. from experimental groups infused with indicated concentrations of *ortho*IL-2. Dotted grey lines indicate mean frequencies of each sub-population across all experimental groups in P14 IL2Rβ-ortho^YFP-^ control cells. E-Experimental design. F-Ratio of cell number between co-transferred P14 IL2Rβ-ortho^YFP+^/P14 Empty^YFP+^ in indicated experimental groups at d35p.i. Combo stands for αPD-L1+*ortho*IL-2 (100KIU). G-Relative frequency of P14 Empty^YFP+^ and P14 IL2Rβ-ortho^YFP+^ cells in indicated experimental groups at d35p.i. H-Representative dot plots of Ly108/CD69-defined subsets among P14 Empty^YFP+^ and P14 IL2Rβ-ortho^YFP+^ cells isolated at d35p.i. from indicated experimental groups. I-Ratio of absolute cell number between indicated subsets of co-transferred P14 IL2Rβ-ortho^YFP+^ and P14 Empty^YFP+^ isolated at d35p.i. from indicated experimental groups.

Because expansion of T_EX_^prog2^ and T_EX_^int^ cells is also observed following PD-1/PD-L1 pathway blockade,^27^ we next tested the potential of PD-1 pathway blockade to combine with *ortho*IL-2. P14 Empty and P14 IL2Rβ-ortho cells expanded similarly upon PD-L1 blockade. However, the P14 IL2Rβ-ortho cells substantially outnumbered their P14 Empty counterpart in the same mice when *ortho*IL-2 was provided at the time of PD-L1 blockade (**Fig. 6F-G** and **S7J-K**). This burst in P14 IL2Rβ-ortho cell number was due to a selective amplification of T_EX_^prog2^ and an even more robust increase in T_EX_^int^ cells (**Fig. 6H-I**). Thus, the strong combinatorial potential of IL-2-derived signals to synergize with PD-1/PD-L1 blockade reported previously^48^ resides in the convergence of the two approaches at amplifying the T_EX_^int^ subset likely in a Stat5-dependent manner.

### Temporal reactivation of Stat5 in T**_EX_** progenitors enables functional recovery and partial epigenetic rewiring towards the T_EFF_/T_MEM_ lineage upon rechallenge

Given the ability of Stat5-dependent signals to restrain exhaustion and foster effector-like biology, we next tested whether engaging Stat5 in combination with strong re-differentiation signals could rewire the epigenetic program of mature T_EX_ cells. To test this idea, we sort-purified RV transduced IL2Rβ-ortho Ly108^+^ T_EX_ progenitors cells on d27p.i., after exhaustion was fully established (**Fig. S8A**). These cells were adoptively transferred into congenic naïve recipient mice and these mice were subsequently challenged with LCMV Arm to provide a strong (re)differentiation signal (**Fig. 7A**). On day 3-to-7 post challenge (p.ch.), recipient mice received daily injections of PBS or *ortho*IL-2 (150KIU) with or without anti-PD-L1 (**Fig. 7A**). We compared these responses to recall responses of conventional memory P14 CD8 T cells (Memory; T_MEM_) isolated from LCMV Arm mice (d>90p.i.) (**Fig. 7A** and **S8A**). When compared head-to-head, T_MEM_ cells numerically outperformed Ly108^+^ T_EX_ progenitors from the PBS-treated group (T_EX_[PBS]) by ∼54-fold on d8 p.ch. (**Fig. 7B**), consistent with previous observations.^11^ T_EX_^[PBS]^ also remained poor cytokine producers, had lower expression of cytolytic molecules (i.e. Gzmb, Gzma), rapidly re-expressed Tox and PD-1, and generated few KLRG1^+^CD127^-^ secondary T_EFF_ compared to the donor T_MEM_ (**Fig. 7C-E**). In contrast, however, IL2Rβ-ortho Ly108^+^ T_EX_ progenitors treated with *ortho*IL-2 (T_EX_^[oIL2]^) during re-challenge underwent robust secondary expansion compared to their PBS-treated counterparts, approaching the expansion potential of T_MEM_ cells, especially when *ortho*IL-2 was combined with PD-L1 blockade (**Fig. 7B**). The accumulation advantage of T_EX_^[oIL2]^ cells versus T_EX_^[PBS]^ persisted at d40 p.ch. (**Fig. S8B**). *Ortho*IL-2 treatment was sufficient to restore polyfunctionality as assessed by IFNγ and TNF production and this polyfunctionality was not further enhanced by addition of PD-L1 blockade (**Fig. 7C**). *Ortho*IL-2 treatment also resulted in higher expression of effector related molecules reaching levels similar to (T-bet, CD94, GzmB) or even higher (GzmA) than observed for secondary T_EFF_ responses derived from T_MEM_ and those qualitative changes occurred in the absence of an increase in the KLRG1^+^ population (**Fig. 7D, E**). Tox expression was reduced in T_EX_^[oIL2]^ cells whereas PD-1 expression remained unchanged compared to the PBS treatment group (**Fig. 7D**), consistent with the additional benefit of blocking the PD-1 pathway in combination with *ortho*IL-2 treatment (**Fig. 7B and D**). Thus, in this rechallenge setting, *ortho*IL-2 treatment synergized with PD-L1 blockade for robust expansion of T_EX_ progenitor cells and accessing the IL-2-STAT5 axis in this setting had a selective qualitative impact on restoring robust expansion, polyfunctionality and effector biology.

**Figure 7:**
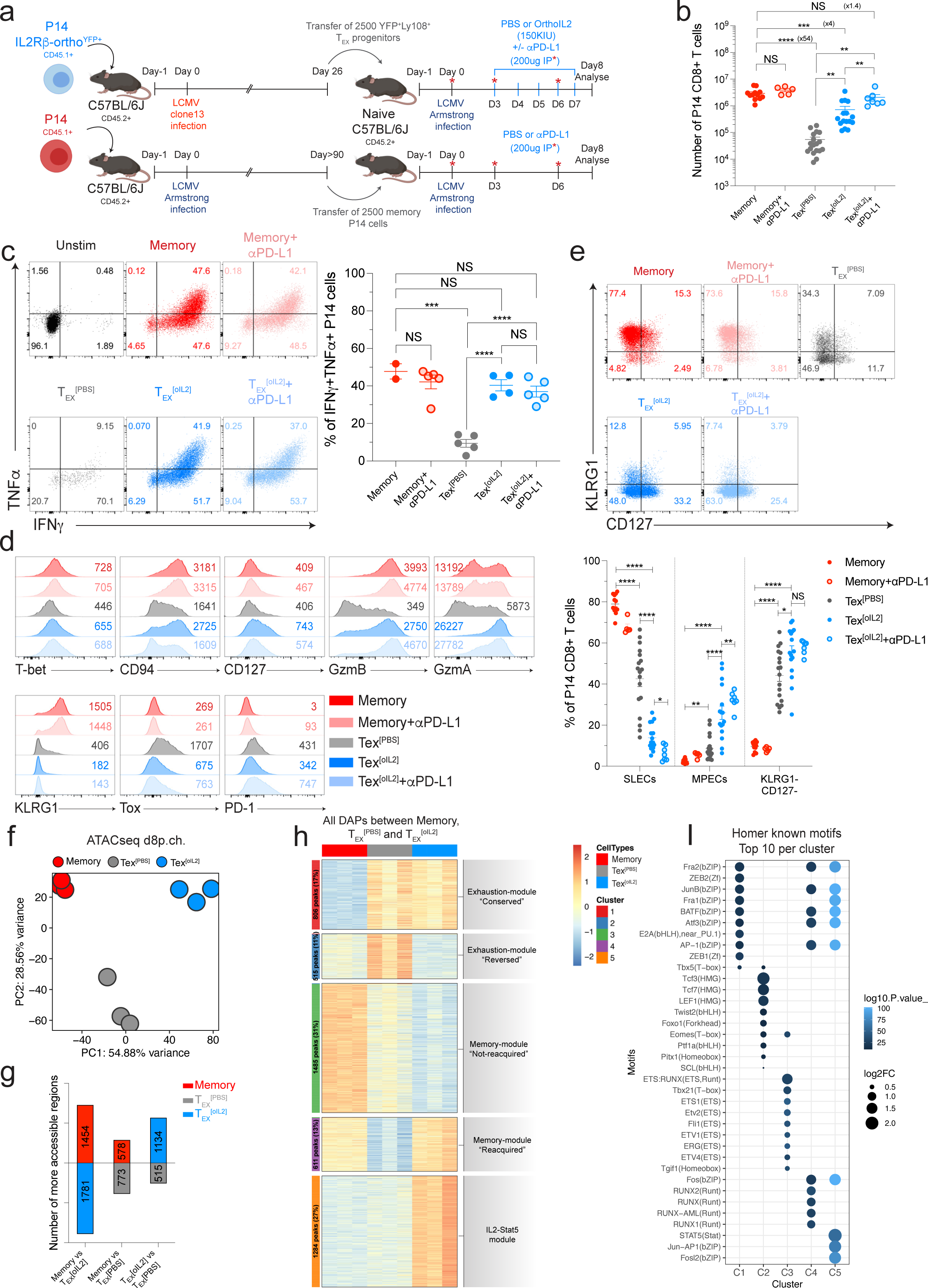
Improved function and partial epigenetic rewiring of rechallenged T_EX_^prog^ cells with targeted IL-2-Stat5 signals. **A**-Experimental design. P14 Memory (Memory) and P14 Ly108^+^ T_EX_ progenitors (YFP^+^, expressing the IL2Rβ-ortho receptor, [T_EX_]) were sorted from indicated time post Arm (d≥90p.i.) or Cl13 (d26p.i.) infection respectively (see **Fig. S8A** for sorting strategy), transferred into new hosts and challenged with LCMV Arm. Mice injected with T_EX_ cells (Ly108^+^ P14 expressing IL2Rβ-ortho^YFP+^) were treated with either PBS (T_EX_^[PBS]^) or daily infusion of *ortho*IL-2 (150KIU day 3-7; [T_EX_^[oIL2]^) in combination or not with αPD-L1 blockade (day0, -3 and -6p.ch.). P14 memory cells were treated with PBS or αPD-L1 at similar time points. Cells were analyzed in the spleen at d8p.ch. **B**-Absolute numbers in the spleen at d8.p.ch. **C**-Cytokine secretion by re-challenged memory and T_EX_ from each experimental conditions after 5h of *in vitro* re-stimulation with gp33 peptide. **D**-Expression of indicated markers on re-challenged memory and T_EX_ from each experimental condition. **E**-Frequency of KLRG1/CD127-defined sub-populations among re-challenged memory and T_EX_ from indicated experimental groups. **F**-PCA of ATACseq data using all DAPs (FDR 0.01, lfc≥2) between indicated populations isolated at d8p.ch. **G**-Number of peaks more accessible in indicated populations and comparisons (FDR 0.01, lfc≥2). **H**-Clustered heatmap (k-means) plotting all DAPs between indicated populations (FDR 0.01, lfc≥2). **I**-Motif enrichment analysis (Homer) plotting the top 10 motifs enriched in DAPs from corresponding clusters in Fig.7H.

To interrogate the mechanisms of this *ortho*IL-2 treatment benefit on T_EX_ cells, we performed ATAC-seq on T_MEM_, T_EX_^[PBS]^ and T_EX_^[oIL2]^ at d8p.ch. Principle component analysis revealed distinct chromatin landscapes for T_MEM_, T_EX_^[PBS]^ and T_EX_^[oIL2]^ with 4701 DAPs (lfc>2, FDR≤0.01) by pairwise comparisons (**Fig. 7F,G, Table S6**). K-means clustering of all DAPs identified modules of open chromatin regions preferentially accessible in cells originating from T_EX_^prog^ (open in T_EX_^[PBS]^ vs T_MEM_; C1 and C2; Exhaustion-Modules) or T_MEM_ cells (open in T_MEM_ vs T_EX_^[PBS]^; C3 and C4; Memory-Modules) (**Fig. 7H, Table S6**). These data indicated that even in settings of strong (de)differentiation signals, scars of the exhaustion epigenetic landscape persisted in cells that expanded from T_EX_^prog^.^11, 13^ Targeted delivery of *ortho*IL-2-signals, however, reversed parts of this epigenetic program in T_EX_^prog^-derived cells (**Fig. 7H**; C2; Exhaustion-module “Reversed”) and even allowed for acquisition of open chromatin patterns associated with the T_EFF_/T_MEM_ lineage (**Fig. 7H**; C4; Memory module “Reacquired”). In addition, *ortho*IL-2 treatment resulted in a selectively increased accessibility at a large fraction of chromatin regions that were otherwise closed in both T_EX_^[PBS]^ and T_MEM_-derived cells (**Fig. 7H**; C5; “IL2-Stat5 module”). Notably, *ortho*IL-2 treatment increased chromatin accessibility at genes related to cell proliferation (*Cdkn2b*), effector differentiation (*Id2*, *Klrb1c*), IL-2/Stat5 responsiveness (*Il2ra*, *Cish*) and interferon response (*Ifitm1*, *Ifitm3*) (**Table S6**). Nevertheless, a fraction of T_EX_^prog^-(C1; Exhaustion-module “Conserved”) and T_MEM_-(C3; Memory module “Not Re-acquired)-related open chromatin regions were not or were more moderately affected by *ortho*IL-2 treatment suggesting selectivity in the epigenetic changes triggered by the *ortho*IL-2-Stat5 axis (**Fig. 7H**). Finally, the genomic regions remodeled in *ortho*IL-2 treated T_EX_^prog^ (T_EX_^[oIL2]^), in particular the IL2-Stat5 C5 module also displayed consistent directionality of chromatin accessibility in P14 STAT5CA cells from d8p.i. (**Fig. S8C**). The susceptibility of those regions to IL-2/Stat5-mediated chromatin accessibility modulation at either early or late time-points of a chronic viral infections suggested opportunities to leverage this IL-2/Stat5 axis for either the prevention or therapeutic reprogramming of T_EX_ cells.

To examine the transcriptional circuitry that was rewired by *ortho*IL-2 signals in this setting of T_EX_^prog^ (re)differentiation, we next compared the network of TF binding site in the altered chromatin accessibility landscape of T_MEM_, T_EX_^[PBS]^ and T_EX_^[oIL2]^. The open chromatin landscape of T_EX_^[PBS]^ enriched in binding motifs for TCR-inducible bZIP domain-containing AP-1 family members ([C1]; i.e. Fra1/2, JunB, BATF, Atf3 or AP-1) and High Mobility Group-TFs ([C2]; i.e. Tcf7, Tcf3, the Tcf7-homologue Lef1 and the Tcf7 partners Foxo1 and Eomes)^49–51^ (**Fig. 7I**). T_MEM_-derived cells, in contrast, enriched for T-box (i.e. Tbx21), ETS (i.e. Ets1) and Runt (i.e. RUNX1/2) motifs (C3 and C4), consistent with distinct transcriptional circuitry governing the T_EX_ and T_MEM_ lineages during recall responses.^25, 26, 40, 52^ Notably, whereas T_MEM_ also contained accessible bZIP motifs, these motifs were located in chromatin accessible regions associated with C4 DAP, whereas the bZIP motifs enriched in T_EX_^[PBS]^ were found mainly in C1 open chromatin regions. These data suggested distinct wiring of TCR-dependent signals (i.e. mediated via bZIP TFs) in T_EX_ versus T_MEM_ during rechallenge. *Ortho*IL-2 treatment did not alter accessibility at bZIP motifs in C1, the module associated with T_EX_^[PBS]^ recall responses. However, *ortho*IL-2 increased accessibility at bZIP motifs in the T_EFF_/T_MEM_-related C4 and also provoked increased accessibility at bZIP binding sites in C5, the module preferentially enriched in the T_EX_^[oIL2]^ cells (i.e. Jun-AP1, Fosl2) (**Fig. 7I**). Thus, *ortho*IL-2, likely through Stat5 engagement, appears to have a prominent impact in shaping the set of bZIP family TF binding sites in T_EX_ during (re)differentiation. Moreover, *ortho*IL-2 treatment also selectively reversed the T_EX_-associated accessibility at HMG-TFs bound regions (C2) and re-engaged T_MEM_/T_EFF_-related enhancers such as those bound by the Runx-family of TFs (C4) (**Fig. 7I**). Together, these data suggest that providing strong (re)differentiation signals via antigenic restimulation in combination with IL-2 and/or Stat5 signals may have therapeutic potential to rewire T_EX_. This augmented IL-2-Stat5-signaling during (re)differentiation of T_EX_^prog^ resulted in a remodeled epigenetic landscape and subsequent reshaping of the TF network in these cells towards a hybrid T_EFF_/T_EX_ state that combined partial silencing and rewiring of exhaustion-related open chromatin regions to re-engagement of some chromatin accessibility regions associated with the T_EFF_/T_MEM_ lineage.

## Discussion

Reversing or rewiring the epigenetic program of T_EX_ remains a major goal of cancer immunotherapy.^11, 13, 25, 53^ Here, we discovered a reciprocal circuit between Stat5a and Tox in which Stat5a antagonizes Tox and the Tox-driven T_EX_ epigenetic program, fostering the acquisition of effector-like biology. In established T_EX_, boosting Stat5 activity partially rewired the T_EX_ open chromatin landscape towards the T_EFF_/T_MEM_ lineage with a preferential ability to function at the point of developmental flexibility that occurs as T_EX_^prog^ convert to the “effector-like” T_EX_^int^ subset. The use of an orthogonal IL-2/IL2Rβ-pair system^47^ allowed Stat5-signals to be directed exclusively to the Ag-specific CD8 T cells of interest *in vivo* and strongly synergized with PD-1 pathway blockade through coordinated expansion of T_EX_^int^ cells. These data may help explain the therapeutic benefit of IL-2 in settings of T cell exhaustion,^54, 55^ the combinatorial effect of IL-2 treatment with PD-1 pathway blockade,^48^ and define mechanisms by which γ_c_-cytokine signaling can impact CD8 T cell exhaustion. Moreover, a notable feature of manipulating Stat5 activity in T_EX_ was the generation of a highly durable hybrid state of differentiation that has features of effector biology, NK receptor expression, resistance to exhaustion, and durability which together, could have considerable therapeutic benefit.

IL-2 was one of the first effective immunotherapies for cancer^55^ and, can have a direct impact on T_EX_.^56^ Our data now provide mechanistic explanations for these effects of IL-2. First, Stat5 antagonizes Tox and the Tox-dependent T_EX_ epigenetic imprinting fostering effector-like differentiation. This Stat5 antagonism of Tox may explain the preferential impact of early Stat5-signals in settings where Tox is abundant (chronic infections, cancer) versus those that favor T_EFF_ (e.g. acutely resolving infections) where Tox expression is low.^7^ Second, Stat5 was necessary for formation of T_EX_^int^ cells a finding that may explain the strong synergy of IL-2 and PD-1 blockade. Mechanistically, Stat5 attenuated or extinguished the stem-like biology of T_EX_^prog^ to initiate exit from quiescence, cell-cycle re-entry and allow downstream T_EX_^int^ cell differentiation. One potential link between these events may be the mechanisms of downregulation of Tcf1 which is essential for exit from the T_EX_^prog^ state.^27, 35, 57^ In other settings, IL-2/Stat5-signals can repress Tcf1 activity and promote cellular differentiation.^58, 59^ Indeed, here we found that enhancing Stat5 activity (STAT5CA) depleted Tcf1^+^ cells and provoked a loss of Tcf1 binding sites in established T_EX_ (*ortho*IL-2) whereas Stat5-deficiency trapped T_EX_ at the progenitor stage. Thus, the balance between Tcf1 and Stat5 activity may be one key regulator node for differentiation of T_EX_^prog^ into downstream T_EX_ subsets including T_EX_^int^. Third, Stat5 promotes the effector circuitry in T_EX_^int^ cells including driving expression of many effector (e.g. *GzmB*, *IFNγ*, *FasL or perforin*) and NK-related genes also previously linked to Stat5 activity in other settings.^38, 58, 60^ Hence, Stat5 not only functions to drive formation of T_EX_^int^ cells but also likely controls expression of some of the key genes associated with this effector/NK-like biology. Together, these observations provide rationale for developing therapeutic strategies to increase Stat5 activity in T_EX_ in settings of chronic infection or cancer. In particular, the *ortho*IL-2 approach^47^ may control for previous limitations by delivering Stat5 inducing signals only to the cells of interest.^61^

Since the discovery of the distinct epigenetic wiring of T_EX_ that limits re-differentiation upon PD-1 blockade,^25, 40^ developing approaches to reprogram the epigenetics of T_EX_ has been a major goal. Identifying such strategies, however, has proven challenging. The data presented here reveal new potential opportunities for, at least partial epigenetic re-wiring of T_EX_. T_EX_ can retain epigenetic “scars” in settings of disease cure and rapidly re-engage the T_EX_ program upon antigenic rechallenge.^11, 13,8,62^ In settings of an acute viral rechallenge, we found that boosting the IL-2/Stat5 signals reversed a substantial fraction of these exhaustion-associated scars and restored accessibility at open chromatin regions associated with the T_MEM_/T_EFF_ lineage. This partial epigenetic reprograming was sufficient to restore robust re-expansion and polyfunctionality. The exhaustion-specific open chromatin regions reversed by the IL-2/Stat5 axis were enriched for HMG-motifs especially those that could be bound by Tcf1. Tcf1 functions in activated CD8 T cells to maintain stemness at the expense of effector differentiation^49, 50, 57, 63^. One possibility is that Tcf1 may restrain T_EFF_ features in T_EX_ and the ability of Stat5 signals to repress Tcf1 activity^58, 59^ maybe be sufficient to relieve the Tcf-mediated T_EX_^prog^ restraint. Coupled to an antagonism of Tox, IL-2/Stat5 signals are likely to foster T_EX_ rewiring by both augmenting a developmental biology conversion of T_EX_^prog^ into T_EX_^int^ by antagonizing Tcf1 and also by removing the Tox-dependent reinforcement of the T_EX_ program. Thus, appropriately accessing the IL-2/Stat5 pathway provides a strong combination of signals for T_EX_ (re)differentiation.

Long-term persistence in settings of continued TCR signaling is a hallmark of T_EX_ cells compared to T_EFF_ or T_MEM_.^14, 15, 35, 64, 65^ Thus, a notable feature of constitutive Stat5a activity in Ag-specific CD8 T cells during chronic infection was the durability of this population despite the relative absence of the key regulators of T_EX_ persistence, Tox and Tcf1.^6-9,31–33, 66^ Although there are some data suggesting a role for IL-2-signals in fine-tuning memory CD8 T cell formation,^67^ IL-2 signals also drive terminal differentiation of short-lived effector CD8 T cells and prolonged exposure to exogenous IL-2 exacerbates T_EFF_ contraction in settings of acute viral infection^54, 68–72^. Moreover, use of IL-2 for *in vitro* expansion in settings of adoptive cell therapy (ACT) has been associated with poor engraftment and/or limited durability or anti-tumor activity of Ag-specific CD8 T cells.^59^ Thus, although IL-2 fosters strong effector function, this cytokine can also drive terminal differentiation.^73, 74^ As a result, in settings of ACT, strategies to temper Stat5-signals (e.g. using engineered IL-2 variants or alternate γ_c_-cytokines during *in vitro* expansion)^58, 59, 75^ have been developed to restrain terminal differentiation and support formation of a stem-like compartment with superior engraftment potential and anti-tumor activity.^36, 58, 59, 68, 69, 76, 77^ Thus, our data on the durability benefits of STAT5CA in chronic viral infection suggest several possibilities. First, constitutively active Stat5 may function differently than prolonged exposure to IL-2. Second, enforcing Stat5-signals directly in CD8 T cells may differ from exogenous IL-2 treatment, especially in settings where the ability of T_EFF_ signal downstream of IL-2 is reduced due to changes in receptor expression and/or signaling efficiency.^36, 78, 79^ Third, continuous IL-2/Stat5 signals may provoke different effects than short-term IL-2 exposure as used in ACT protocols^59^ or previous studies only providing additional IL-2 during the effector phase.^54^ However, one last possibility is that in the setting of continuous TCR signals that drive exhaustion, enforced Stat5 activity synergizes with other exhaustion-driven antigen-dependent survival signals. Dissecting these questions will be an important future goal to determine how Stat5 interacts with other signals and devise strategies to optimally exploit the Stat5/IL-2 pathway for enhancing immunotherapy. In summary, we identify a role for augmented IL-2/Stat5 signals in a potential epigenetic rewiring of T_EX_ cells and uncover the underlying molecular and cellular mechanisms for these effects. The result of increasing IL-2/Stat5 signals is a hybrid differentiation state combining therapeutically useful features of both T_EFF_ and T_EX_ leading to improved control of disease. The use of the *ortho*IL-2 system demonstrated that these effects are cell intrinsic to T_EX_ and suggests future strategies for Stat5 targeting therapeutics including cytokine-based and engineered cellular therapy-based approaches. Future studies in humans will be necessary to understand how these molecular principles extend to more complex settings with both pre-existing T_EX_ and opportunities for new T cell priming as well as role for other γ_c_ responsive cell types. Nevertheless, these data may provide a guide for developing and evaluating such therapies in future clinical trials.

## STAR★METHODS

### Mice

Six-week old C57BL/6 female mice (CD45.2, Charles River, NCI) were used as recipient mice for most adoptive transfer experiments. Alternatively, six-week old C57BL/6 male or female (CD45.2, The Jackson Laboratory) mice were used as recipients for Stat5iKO experiments. P14 TCR transgenic mice expressing a TCR specific for the LCMV D^b^gp33-41 peptide were bred in house and backcrossed onto the C57BL/6 background. P14 Rosa^YFP^ *Stat5a/b*^fl/fl^ (P14 Stat5iKO) mice were generated by crossing *Stat5a/b*^fl/fl^ mice (The Jackson laboratory, ref-#032053-JAX) with P14 Rosa^YFP^ mice (bred in house). All experiments and breeding conditions were in accordance with Institutional Animal Care and Use Committee (IACUC) guidelines for the University of Pennsylvania.

### Viruses and Infections

LCMV Arm and Cl13 were grown in BHK cells and titrated using plaque assay on VERO cells as described.^80^ Recipient mice were infected either intraperitoneally (i.p.) with LCMV Arm (2×10^5^ plaque forming units [PFU]) or intravenously (i.v.) with LCMV Cl13 (4×10^6^ PFU).

### Cell line and tumor transplant

The B16_gp33_ melanoma cell line was maintained in DMEM supplemented with 10% FBS, 1% L-glut and 1% Pen/Strep. Tumor cells cultured for less than two weeks were resuspended in cold PBS and implanted subcutaneously (5×10^5^ cells in 50μl) in the flank of recipient mice using 29G1/2 syringes. Tumor size was monitored every two days using a digital caliper and mice were euthanized before tumors exceeded the volume permitted by the IACUC guidelines for the University of Pennsylvania.

### Retroviral vectors

The STAT5CA construct has been described previously^36, 37^ and was kindly provided by Dr. Susan Kaech (The Salk Institute). The IL2Rβ-ortho construct has been described previously^47^ and was obtained from Dr. Christopher K. Garcia (Stanford University) under the Material Transfer Agreement RIS#59882/00 between Stanford University, the University of Pennsylvania and the Parker Institute for Cancer Immunotherapy (PICI). Both constructs were cloned into a MSCV-IRES plasmid containing either VEX or YFP/GFP-reporters. RV particles were produced by transfection of 293T cells. Briefly, 293T cells were pre-incubated with warmed cDMEM supplemented with chloroquine (25μM; Sigma). Cells were transduced with a pCL-Eco plasmid (15μg) and MSCV-IRES expression plasmid (15μg) using Lipofectamine 3000 (ThermoFisher Scientific) for 6 hours at 37°C 5%CO2. After incubation, transduction medium was replaced with fresh cDMEM. RV supernatants were collected at days 3 and 4 of culture and titrated on NIH3T3 cells.

## METHOD DETAILS

### Adoptive T cell transfer

PBMCs containing 1×10^3^ P14 CD8 T cells were adoptively transferred into recipient mice 24h prior to infection with either LCMV Arm or LCMV Cl13. For Stat5iKO experiments, P14 Rosa^YFP+/-^ *Stat5a/b*^fl/fl^ (P14 Stat5iKO) and their control counterpart P14 Rosa^YFP+/-^*Stat5a/b*^+/+^ (both CD45.1.2^+^) were harvested from PBMCs and cultured in serum free RPMI medium containing (cre+) or not (cre-) 50μg/ml of tat-cre recombinase (Proteomic Core Facility-Children’s Hospital of Philadelphia) for 45min at 37°C, 5%CO2. Cells were washed once in FBS then complete RPMI (cRPMI), resuspended in cold PBS and 1.5×10^3^ of each were adoptively transferred into separate naïve CD45.2 recipients 24h before infection.^27^ Markers associated with early T cell activation (i.e. CD69, Ly6C, PD-1, CD25, CD62L, CD127) were assessed in P14 populations before infusion into recipient mice to ensure transfer of phenotypically naïve T cells.

### Retroviral (RV) transduction

RV transduction of P14 CD8 T cells was performed as described previously ^81^ with slight modifications. For each experiment, P14 CD8 T cells were enriched from spleens of P14 transgenic mice using EasySep^tm^ CD8^+^ T cell isolation Kit (StemCell) and activated *in vitro* in cRPMI supplemented with αCD3 (1μg/ml), αCD28 (0.5μg/ml) antibodies and rhIL-2 (100U/ml) (PeproTech) at a seeding density of 1×10^6^ cells/ml. One day post activation (between 24-27h), CD8 T cells were re-suspended at a density of 3-5×10^6^ cells/ml mixed with RV supernatant containing polybrene (4μg/ml) at a 1:1 ratio (v/v) and spin-transduced 75’ at 2000g 32°C. After transduction, 4ml of warmed cRPMI containing αCD3, αCD28 and rhIL2 was gently added to each well of a 6-well plate for a final volume of 6ml. Cells were incubated O/N (∼16h) at 37°C, 5% CO2. The next day, transduced cells were stained for 15min with LiveDead Aqua (ThermoFisher Scientific) or Zombie NIR (BioLegend) and anti-CD8 antibodies in 1XPBS at RT, resuspended in warmed cRPMI and RV-positive cells (either VEX+ of YFP/GFP+) were sorted among live CD8 T cells (LiveDead Aqua/Zombie NIR^-^CD8^+^). For most RV experiments described, P14 cells expressing different congenic markers (CD45.1 or CD45.1.2) were used for transduction of control RVs (empty) and RVs encoding proteins of interest (STAT5CA or IL2Rβ-ortho). The two congenically distinct P14 populations were then mixed at a 1:1 ratio in warmed PBS and injected into C57BL/6 recipients (2.5×10^4^ each) infected 3 days earlier with LCMV Arm or Cl13.

### Tumor experiments

C57BL/6 mice were inoculated with 5×10^5^ B16_gp33_ cells. Ten days post tumor inoculation, mice were randomized and either left untreated or injected i.v. with FACs purified P14 Empty or P14 STAT5CA cells (5×10^5^).

### Cell preparation, flow cytometry and cell sorting

Spleens were mechanically disrupted onto a 70μM cell strainer using the plunger of a 3mL syringe and resuspended in 1mL of ACK red blood cell lysing buffer (Gibco) for 3 min at room temperature (RT). Cell suspensions were washed and resuspended in cRPMI supplemented with 10% FBS, 1% penn/strep, 1% L-glut, Hepes 10mM (Cell Center, UPenn), MEM non-essential amino acids 1% (Gibco), Sodium Pyruvate 1mM (Cell Center UPenn), β-mercaptoethanol (0.05mM). Bone marrow suspensions were harvested by flushing cells out of the femur and tibia of infected mice with a 29G syringe and cRPMI. Cells were then treated as above. For lungs and livers, mice were perfused with cold PBS. Lungs were cut in a petri dish, disrupted in 10 ml of RPMI (1%FBS) in the presence of Collagenase D (1X) (Roche) using a MACs dissociator (Miltenyi Biotec) and incubated for 45min at 37°C under agitation. After incubation, lung cells were disrupted a second time on a MACs dissociator (Miltenyi Biotec) and processed as above. After mechanical disruption onto a 70μM strainer, lymphocytes from livers were enriched using Percoll (GE Healthcare) density gradient separation (80%/40%), washed two times with cRPMI and processed as above. Blood samples were collected in 1ml of PBS 2mM EDTA. RPMI was added (1ml) and samples were underlaid with 1ml of Histopaque 1083 (Sigma Aldrich) for lymphocyte enrichment using density gradient concentration. Remaining red blood cells were lysed using ACK lysing buffer (Gibco) for 3min at RT. Equal numbers of cell were stained with extracellular antibodies for 30min on ice in FACs buffer (PBS 1X, 1% FBS, 2mM EDTA) in the presence of Live/Dead Fixable Aqua Cell Stain (ThermoFisher Scientific). Cells were then fixed for 20 min on ice with Cytofix/Cytoperm (BD bioscience) and analyzed by flow cytometry. For cytoplasmic protein detection, cells were incubated for an additional 30min on ice in Perm/Wash buffer (BD bioscience) and stained for 1h on ice in Perm/Wash buffer (BD bioscience) containing antibodies targeting cytoplasmic proteins (active-caspase3, gzmA, gzmB, IFNγ, TNF). For TFs detection, cells were fixed (20min) and permeabilized (30min) on ice using the Foxp3 Transcription Factor buffer set (ThermoFisher Scientific) and incubated for an hour with TF antibodies. For TFs detection in cells expressing a fluorescent reporter protein (VEX or GFP/YFP), cells were pre-fixed 5min in 2% formaldehyde (ThermoFisher Scientific) before fixation and permeabilization using the Foxp3 TF buffer set (ThermoFisher Scientific). Samples were resuspended in FACs buffer, acquired on an LSR II or BD FACSymphony and analysed with FlowJo v.10 software (Tree Star Inc).

For cell sorting *ex vivo*, CD8 T cells were enriched from total splenocytes using the EasySep^tm^ CD8^+^ T cell isolation Kit (StemCell) (routinely >90% purity), stained on ice for 30’ with relevant cocktails of antibodies and populations of interest were sorted at 4°C on an BD FACSARIA (BD Bioscience) using a 70 μM nozzle in 50% FBS RPMI. Purity was routinely >95%. For ATACseq, scRNAseq and CITEseq experiments, RV-positive or reporter expressing P14 cells (either VEX^+^ or GFP/YFP^+^) were sorted among LiveDead Aqua/ZombieNIR^-^CD8^+^CD45.1^+^ cells. For re-challenge experiments, memory and T_EX_^prog^ P14 cells were sorted among LiveDead Aqua^-^CD45.1^+^CD45.2^-^CD8^+^ T cells and T_EX_^prog^ cells were further discriminated as Ly108^+^Cx3cr1^-^.

### Intracellular cytokine staining

Splenocytes or total CD8 T cells enriched using the EasySep^tm^ CD8^+^ T cell isolation Kit (StemCell) (1-2×10^6^) were re-stimulated *in vitro* for 5h at 37°C 5% CO2 in cRPMI supplemented with GolgiStop (1/250; BD bioscience), GolgiPlug (1/500; BD bioscience) and gp_33-41_ peptide (NIH, 0.4μg/ml). Cells were then washed and stained using the BD Fixation/permeabilization kit (BD Bioscience).

### Antibody and cytokine treatment

Where indicated, mice were depleted of CD4 T cells using two i.p. injections of 200μL of PBS containing 200μg of monoclonal anti-CD4 antibody (clone GK1.5, BioXcell) one day prior and post infection with LCMV Cl13. PD-L1 blockade was performed in CD4-depleted mice as previously described.^25^ Sequential i.p. injections of 200μl of PBS containing or not rat anti-mouse PD-L1 monoclonal antibody (200μg/injection, clone 10F.9G2, BioXcell) were performed every three days between days 22 and 34 for a total of five injections. For re-challenge experiments, similar injections were performed at d0, 3 and 6 post infection with LCMV Arm. For experiments using the IL2/IL2Rβ-orthogonal pair system, *ortho*IL-2 was infused daily (I.P) in 200μl of cold PBS at indicated concentrations from d21-to-25p.i. In some experiments, groups of mice were treated similarly with regular mIL2 as a reference (25KIU/injection). In experiments combining *ortho*IL-2 treatment with PD-L1 blockade, *ortho*IL-2 was infused I.P every 2 days (100KIU/injection) for the duration of PD-L1 treatment (d22-34p.i.). For re-challenge experiments, *ortho*IL-2 was infused daily (I.P) from d3-to-d7 post challenge (150KIU/injection).

### Active caspase-3 and BrdU detection

Splenocytes from infected mice were incubated for 5 hours at 37°C 5%CO2 in cRPMI prior intra-cytoplasmic detection of active-caspase 3 (BD Bioscience) using BD Fixation/Permeabilization kit (BD Bioscience). Mice adoptively transferred with either P14WT or P14Stat5iKO were injected I.P with 2mg of BrdU at d7p.i. with LCMV Cl13 and BrdU detection in splenic P14 cells was performed one day later (d8p.i.) using a BrdU detection Kit (BD Bioscience) according to manufacturer’s protocol.

### Sample preparation for Cut&Run

Cut&Run was performed as previously described ^82^ with modifications. P14 Empty and P14 STAT5CA cells were sorted at d8.p.i. with either LCMV Arm or Cl13 from recipients of two independent experiments and 0.5 to 3×10^5^ cells were recovered in 1.5ml DNA LoBind Eppendorfs containing 650μl of 50%FBS RPMI. Samples were washed twice in 1ml of cold wash buffer (20mM HEPES-NaOH pH7.5, 150mM NaCl, 0.5mM Spermidine and protease inhibitor from Roche), re-suspended in 400μl (final) of wash buffer containing 20μl of BioMagplus Concanavalin A-coated magnetic beads (Bangs Laboratories) per reaction and rotated for 15min at 4°C to allow the cells to bind. Tubes were placed on a magnet stand and liquid was removed. Beads were then incubated O/N at 4°C in 250μl of antibody buffer (20mM HEPES-NaOH pH7.5, 150mM NaCl, 0.5mM Spermidine, 2mM EDTA, 0.1% digitonin and protease inhibitor from Roche) containing 2.5μl (1/100) of antibodies against H3K27ac (Active Motif) or IgG control (Cell Signalling Tech). Samples were then washed twice in 500μl of Digitonin Buffer (20mM HEPES-NaOH pH7.5, 150mM NaCl, 0.5mM Spermidine, 0.1% digitonin and protease inhibitor from Roche), resuspended in 250μl of cold Digitonin Buffer containing Protein-A micrococcal nuclease (pA-MN) and rotated at 4°C for 1h. Beads were washed twice in 1ml of cold Digitonin Buffer to remove unbound pA-MN, resuspended in 150μl of Digitonin Buffer, cooled down at 0°C on a pre-cooled metal block for 5min and incubated 30min at 0°C with CaCl_2_ (3μl of 0.1M per sample) to initiate pA-MN digestion. Reaction was stopped by addition of 150μl of 2X stop Buffer (340mM NaCl, 20mM EDTA, 4mM EGTA, 0.02% Digitonin, 50μg/ml RNAseA and 50μg/ml Glycogen) followed by 10min incubation at 37°C to release target chromatin. Samples were then centrifuged 5min 16,000g 4°C and supernatants were transferred to new tubes. Chromatin fragments were incubated 10min at 70°C with 3μl of 10% SDS and 2.5μl of proteinase K (20mg/ml) followed by phenol/chloroform/isoamyl alcohol-based extraction according to original protocol (method B). Upper phase containing DNA was mixed with 1μl of glycogen (20mg/ml) and incubated with 750μl of cold 100% ethanol at -20°C O/N. Samples were centrifuged 30min 16,000g 4°C, rinsed once with 1ml of cold 100% ethanol and centrifuged again for 5min 16,000g 4°C to remove residual ethanol. Samples were air-dried, resuspended in 50μl of molecular grade water and stored at -20°C. DNA libraries were built using the NEBNext Ultra II DNA Library Prep Kit for Illumina (NewEngland Biolabs) with the following modifications.^83^ NEBNext End Prep step was performed using 25μl of input material for a final volume of 30μl and the following adapted program (30min-20°C, 60min-50°C, Hold at 4°C). Adaptor was diluted at 1:25 and added at 1.5μl for ligation (15min-20°C) followed by addition of 1.5μl of Red USER Enzyme and additional 15min incubation at 37°C. Size selection was performed using 80μl of AMPure XP beads (Beckman Coulter) and purified DNA fragments were amplified for 14 cycles (annealing time changed to 10s). Libraries were cleaned-up with two rounds of size selection with AMPure XP beads (24μl/12μl; Beckman Coulter) and eluted in 15μl of molecular grade water, and amplicons quality was assessed on a 2200 TapeStation (Agilent Technologies). Libraries were quantified by qPCR using the NEBNext Library Quant kit for Illumina (NewEngland Biolabs) according to manufacturer’s protocol and pooled at equal molarity (1nM). Denatured Libraries were diluted at 1.8pM, loaded into a NextSeq 500/550 High Output Kit (75 cycles, Illumina) and paired-end sequencing was performed on a NextSeq 550 (Illumina).

### Sample preparation for scRNAseq

Splenocytes from recipient mice were pooled from duplicate experiments and CD8 T cell enrichment was performed using EasySep^tm^ CD8+ T cell isolation Kit (StemCell). Enriched CD8 T cells were stained and P14 populations of interest were sorted at 4°C in 1.5ml Eppendorf tubes containing 50% FBS RPMI as described above. Sorted samples were topped with cold PBS 0.04% BSA, centrifuged for 5’ 350g at 4°C, washed two times in cold PBS and resuspended in 50-100μl of cold PBS. Samples were counted, down-sampled and equivalent number of cells (6300) between samples were loaded into the Chip (Chromium Next GEM Chip G) of a Chromium Next GEM Single Cell 3’ Reagent Kit v3.1 (Dual Index, 10x Genomics) and run onto a Chromium Controller. Samples were then processed according to manufacturer’s protocol. cDNA libraries were prepared using the Dual Index TT Set A (10x Genomics) and the number of indexing PCR cycles was adjusted to the cDNA input of each individual sample according to manufacturer’s recommendations. Libraries were quantified by qPCR using a KAPA Library Quant Kit (KAPA Biosystems). Normalized libraries were pooled (2.5nM), loaded onto a NovaSeq 6000 SP Reagent Kit (100 cycles, Illumina) for a final concentration of 450pM and paired-end sequencing was performed on a NovaSeq 6000 (Illumina).

### Sample preparation for CITEseq

CITEseq samples from duplicate experiments were prepared as described above (scRNAseq section) and processed according to the CITE-seq protocol from the New York Center Technology Innovation Lab (https://cite-seq.com/protocols/). Briefly, enriched CD8 T cells were incubated for 10’ at 4°C in Staining buffer (2%BSA/0.01% Tween in PBS) containing FcBlock (1/10 dilution ;TruStain^TM^ FcX, Biolegend) followed by a 30’ incubation in Staining Buffer containing TotalSeqB antibodies against Ly108, CD69, Tim-3, PD-1, CD127, CD122, Lag-3, CD38 and KLRG1 (BioLegend) previously titrated according to manufacturer’s protocol using PE-conjugated version of each antibodies. Samples were then washed, sorted as described above, down-sampled and equivalent number of cells (10^4^) between samples were loaded onto the Chip (Chromium Next GEM Chip G) of a Chromium Next GEM Single Cell 3’ Reagent Kit v3.1 (Dual Index, 10x Genomics) and run onto a Chromium Controller. Samples were then processed according to manufacturer’s protocol. Gene expression and Cell surface Protein libraries were constructed using Dual Index TT Set A and Dual Index NT Set A (10x Genomics) respectively. Libraries were quantified by qPCR using a KAPA Library Quant Kit (KAPA Biosystems). Normalized libraries were pooled (0.23nM), diluted to 1.8pg/ml and loaded onto a NextSeq 500/550 High Output Kit v2.5 (150 cycles, Illumina) and paired-end sequencing was performed on a NextSeq 550 (Illumina).

### Sample preparation for ATACseq

ATACseq sample preparation was performed as described ^41^ with minor modifications. Sorted cells (2-to-5×10^4^) were washed twice in cold PBS and resuspended in 50μl of cold lysis buffer (10mM Tris-HCl, pH 7.4, 10mM NaCl, 3mM MgCl2 and 0.1% IGEPAL CA-630). Lysates were centrifuge (750xg, 10min, 4°C) and nuclei were resuspended in 50μl of transposition reaction mix (TD buffer [25μl], Tn5 Transposase [2.5μl], nuclease-free water [22.5μl]; (Illumina)) and incubated for 30min at 37°C. Transposed DNA fragments were purified using a Qiagen Reaction MiniElute Kit, barcoded with NEXTERA dual indexes (Illumina) and amplified by PCR for 11 cycles using NEBNext High Fidelity 2x PCR Master Mix (New England Biolabs). PCR products were purified using a PCR Purification Kit (Qiagen) and amplified fragments size was verified on a 2200 TapeStation (Agilent Technologies) using High Sensitivity D1000 ScreenTapes (Agilent Technologies). Libraries were quantified by qPCR using a KAPA Library Quant Kit (KAPA Biosystems). Normalized libraries were pooled, diluted to 1.8pM, loaded onto a NextSeq 500/550 High Output Kit v2.5 (150 cycles, Illumina) and paired-end sequencing was performed on a NextSeq 550 (Illumina).

## QUANTIFICATION AND STATISTICAL ANALYSIS

### FlowSOM analysis

Compensated parameters for gated P14 Empty and P14 STAT5CA cells were exported from four individual mice co-transferred with both P14 populations and concatenated. Concatenated files were down-sampled using the FlowJo DownSampleV3 plugin for even representation or P14 Empty and P14 STAT5CA populations (15000 cells each), grouped using the t-sne function of FlowJo V10.8.0 using 12 parameters (CD44, Tbet, Tcf1, Tim-3, GzmB, Tox, Lag3, Icos, Ly108, CD39, CD127 and PD-1) and clusters were defined with the FlowSom plugin using the same parameters.

### Ingenuity Pathways Analysis (IPA)

DEGs between P14 WT and P14 ToxKO (**Fig. S1B**),^7^ or cluster specific DEGs (**Fig. S1C-G**) from reprocessed scRNAseq of WT and ToxKO D^b^gp33^+^ CD8 T cells isolated at d7p.i. with LCMV Cl13 (GEO Accession number: GSE119943)^6^ were used as input to the Upstream regulator analysis part of the Core analysis using QIAGEN’s Ingenuity Pathway Analysis (IPA, QIAGEN Redwood City, www.qiagen.com/ingenuity) to generate Transcription factor specific networks.

### Taiji Rank Analysis

Transcription Factor Binding Site (TFBS) analysis and PageRank analysis were performed using Taiji^34^ (https://taiji-pipeline.github.io/algorithm_PageRank.html) and paired ATACseq and RNAseq datasets of indicated T_EX_ subsets (GEO accession number: GSE149879)^27^ to generate TF ranks visualized as heatmap using R pheatmap package (**Fig. S1H**). For **Fig. 1A**, the fold change in Taiji score for T_EX_^int^ cells compared to other T_EX_ subsets was calculated for each individual TF enriched in both the IPA analysis (**Fig. S1B**) and the Taiji Rank analysis (**Fig. S1H**).

### ATACseq

Raw ATACseq FASTQ files from paired-end sequencing were processed using the script available at the following repository (https://github.com/wherrylab/jogiles_ATAC). Samples were aligned to the GRCm38/mm10 reference genome using Bowtie2. We used samtools to remove unmapped, unpaired, mitochondrial reads and ENCODE blacklist regions were also removed (https://sites.google.com/site/anshulkundaje/projects/blacklists). PCR duplicates were removed using Picard. Peak calling was performed using MACS v2 (FDR q-value 0.01). For each experiment, we combined peaks of all samples to create a union peak list and merged overlapping peaks with BedTools *merge*. The number of reads in each peak was determined using BedTools *coverage.* Differentially accessible peaks were identified following DESeq2 normalization using a lfc of 2 and FDR cut-off <0.01 unless otherwise indicated. DAPs were clustered using the k-means clustering methods and motif enrichment analysis was performed for each cluster on indicated DAPs using Homer (default parameters). For peak tracks representation, bed files for each replicate were imported into the UCSC Genome browser online tool. Replicates for each sample were merged and each biological sample were normalized. For sample distance, a distance matrix was calculated using the “euclidean” measure for all peak and plotted as a heatmap.

### CUT&RUN and ChIP-Seq data processing and analysis

Data qualities were checked using FastQC and MultiQC. Paired-end reads were aligned to mm10 reference genome using Bowtie2 v2.3.5 with options suggested by Skene et *al*. 2018.^82^ Bam files containing uniquely mapped reads were kept using Samtools v1.1. MarkDuplicates command from Picard tools v1.96 was used to remove presumed PCR duplicates. Blacklist regions defined by ENCODE were removed, and filtered typical chromosomes were used for downstream analysis. Read per million (RPM) normalized bigwig files to visualize binding signals were created using deepTools bamCoverage v3.3.2 with parameters --normalizeUsing CPM -bs 5 –smoothLength 20 –skipNAs. Biological replicates were pooled together using bigwigCompare with parameter --operation add -bs 5 --skipNAs. Peaks were called on using MACS v2.1 using the broadPeak setting with general adjusted *p*-value cutoff of 0.05. Genes proximal to peaks were annotated against the mm10 genome using R package rGREAT. Venn diagram of peak comparisons were plotted using Bioconductor package ChIPpeakAnno. Peaks of all conditions were merged to create the final union peaks list. For visualization purpose, bigwig files of biological replicates were pooled using wiggletools with mean setting, and median background were subtracted. Tracks were loaded to UCSC genome browser for visualization. Published ChIP-Seq data were downloaded from NCBI (GEO accession number: GSE64407 [Nfat1], GSE98654 [Nfat2], GSE100674 [Stat5]).^42, 43, 84^ Paired-end reads were aligned to mm10 reference genome using Bowtie2 with same parameters as CUT&RUN. RPM normalized tracks were generated using deepTools bamCoverage. Some downloaded signal track files were lifted from mm9 to mm10 using UCSC tool liftOver. Read counts under peaks were generated using deepTools multiBigwigSummary v3.3.2. Binding motifs enrichment analysis were identified using findMotifsGenome.pl from HOMER v4. Local motif binding positions are identified using FIMO with parameters –bfile –motifs. Dot plots were generated using R packages ggplot2.

### Single-Cell RNA sequencing (scRNAseq)

Sample demultiplexing, alignment, filtering and creation of a UMI count matrix were performed using Cell Ranger software v.4.0.0 (10x Genomics). A Seurat object was created from the UMI count matrix using Seurat_4.0.5.^85^ Cells with fewer than 200 or greater than 2500 detected genes were excluded from downstream analysis as of cells with >10% of mitochondrial gene counts. Genes which expression was detected in 3 cells or less were excluded. A total of 920 P14 Empty and 302 P14 STAT5CA cells passed filters with an average sequencing depth of 1984 genes per cell and were considered for downstream analysis. Counts were normalized by total expression in the corresponding cell using the “LogNormalize” function and default scaling factor of 10,000 to give counts per million. Top 2000 variable features were determined using the “vst” selection method. Linear dimensional reduction (PCA) was performed on scaled variable features and features from the 20 most significant PCs were used as input for unsupervised clustering using the “FindNeighbors” and “FindClusters” functions of Seurat with a resolution of 0.3. We next ran non-linear dimensional reduction (UMAP) to visualize the data. Differentially expressed genes were identified by the Seurat function “FindAllMarkers” with min.pct=0.25 and logfc.threshold=0.25 and the top 10 genes per cluster were used for creating the Heatmap using the R package “Dittoseq” (**Fig. S4F**). For projection of indicated gene signatures (SLEC, Exhaustion, Tox program and T_EX_^prog1^, T_EX_^prog2^, T_EX_^int^, T_EX_^term^), Seurat clusters were used as features to calculate module scores of single cells using the “AddModuleScore” of Seurat_4.0.5. Module scores for each of the gene signatures were used to color the UMAP projection (**Fig. S4G**) or dot plots (**Fig. 4F**). Single-cell analysis of P14 Empty and P14 IL2Rβ-ortho was processed independently using similar pipeline. A total of 925 P14 Empty and 951 P14 IL2Rβ-ortho passed filters with an average sequencing depth of 1786 genes per cell and were considered for downstream analysis. Non-linear dimensional reduction (UMAP) was used to visualize data (**Fig. S7E,F**) from the 12 most significant PCs using a resolution of 0.1.

For projection of Stat5a and Tox signatures, scRNAseq data of P14 CD8 T cells isolated from LCMV Cl13 infected mice at d7 and d30p.i. (GEO accession number: GSE131535 and GSE150370),^11, 35^ were reprocessed and module scores for Stat5a and Tox signature genes were used to color the UMAP (**Fig. 1B**) as described above.

### Cellular Indexing of Transcriptomes and Epitopes by sequencing (CITE-seq)

A UMI count matrix was created for P14 WT and P14 Stat5iKO cells using CellRanger 4.0.0. and the two matrixes were used to create Seurat objects. For each sample, an antibody (“adt”) assay was created and added to its cognate Seurat object (WT or Stat5iKO) that were subsequently merged into one object containing rna and adt counts for each sample. Cells with fewer than 200 or greater than 2500 detected genes were excluded from downstream analysis as of cells with >12% of mitochondrial gene counts. Genes which expression was detected in 3 cells or less were excluded. A total of 4377 P14 WT and 4906 P14 Stat5iKO cells passed filters with an average sequencing depth of 1210 genes per cell and were considered for downstream analysis. Counts (rna assay) were then normalized, and the top 2000 variable features were scaled before running linear dimensional reduction (PCA). The 30 most significant PCs were used as input for unsupervised clustering using the “FindNeighbors” and “FindClusters” functions of Seurat with a resolution of 0.1. We next ran non-linear dimensional reduction (UMAP) using the rna assay to visualize Seurat clusters (**Fig. 5F**) or individual samples (**Fig. 5G**). DEGs were identified by the Seurat function “FindAllMarkers” with min.pct=0.25 and logfc.threshold=0.25 and the top 52 variable features by p.val.adj were used for creating the Heatmap (Fig. 5H). The “FindMarkers” function of Seurat was also used for cluster-wise assessment of the number of DEGs using as an input Seurat clusters (**Fig. S6C**) or oligo-tagged antibodies-defined populations (**Fig. 5I**). These oligo-tagged defined populations were delineated based on Ly108 and CD69 adt values with cut-off for positive and negative cells set up using the “FeatureScatter” function of Seurat. DEGs between oligo-tagged defined populations were presented as Volcano plots (**Fig. 5J**). For projection of indicated gene signatures (T_EX_^prog1^, T_EX_^prog2^, T_EX_^int^, T_EX_^term^), oligo-tagged defined populations were used as features to calculate module scores of single cells using the “AddModuleScore” of Seurat_4.0.5. Module scores for each of the gene signatures were used to color the UMAP projection (**Fig. S6E**) or dot plots (**Fig. S6F**).

### Gene ontology

Gene ontology of gene sets of interest were obtained using the Metascape online tool (http://metascape.org/gp/index.html#/main/step1). Pathway enrichment analysis (GO Biological processes) was set for a minimum overlap of 3, a *p*-value cut-off of 0.01 and a minimum enrichment score of 1.5.

### Statistical analysis and experimental replications

Statistics on flow cytometry data were performed using unpaired or paired (co-adoptive transfer experiments) two-tailed Student’s t test. For data presented as a ratio (**Fig. 2D-E, 4B, 5A, and 6F,I**) a Wilcoxon signed rank test was performed with a hypothetical value of 1 or equal to the mean in control group (**Fig. 6F,I**) (GraphPad Prism v6; *p < 0.0332, **p < 0.0021, ***p < 0.0002, ****p < 0.0001). For statistics on scRNAseq data, a Pearson correlation coefficient was calculated as well as a *p* value of significance to estimate the degree of correlation between Stat5a and Tox signatures (**Fig. 1C** and **S1I**). A Wilcoxon t test was performed in **Fig. 4I** to compare enrichment of indicated signature in T_EX_ clusters.

The experiments described were replicated as follows. **Figure 2** - (**B**) N=3 independent experiments (ind exp) with 12 mice/group (**D**) N=2 (Tcf1) or 4 (Tox) with 8 (Tcf1) or 15 (Tox) mice/group (**E**) N=2-4 ind exp with 6-16 mice/group (**F-G**) N=3 ind exp with 10-12 mice/group (**H**) N=2 with 7-9 mice/group. **Figure 4** - (**A**) Representative of 2 ind exp with 10 mice/group (**B**) N=2-4 ind exp with 5-17 mice per time points (**C**) Representative of 2 ind exp with 9-10 mice/group (**K-L**) Representative of 2 ind exp with at least 6 mice/group in each. **Figure 5** - (**A**) N=2-5 ind exp with 6-18 mice/group (**B**) N=4 ind exp with 14 mice per group (**C**) N=3 ind exp with 9-10 mice/group (**D**) N=1-2 ind exp with 2-8 mice/group (**E**) Representative of 2 ind exp with 8-10 mice per group. **Figure 6** – (**B**) N=2 ind exp with 6-10mice/group (**C**) N=2 with 6 mice/group (**D**) N=2 with 6-17 mice/group (**F-I**) N=2 with 9-15 mice per group. **Figure 7** - (**B**) N=5 with 5-18 mice/group (**C**) N=2 with 2-8 mice/group (**D**) Representative of 5 in exp with 5-18 mice/group (**E**) N=5 with 5-18 mice/group.

## Acknowledgments

We thank all members of the Wherry Lab and Dr. Golnaz Vahedi (UPENN) for insightful discussions and comments on the manuscript as well as Dr. Susan Kaech (Salk Institute for Biological Studies) for providing the original STAT5CA constructs. This work was supported by the Parker Institute for Cancer Immunotherapy (PICI) and the National Institute of Health (NIH) grants AI155577, AI115712, AI117950, AI108545, AI082630 and CA210944 (to EJW). ACH is supported by NIH (K08CA230157) and received Doris Duke Clinical Scientist Development and Damon Runyon Clinical investigator Awards. JC-B is a Parker Institute for Cancer Immunotherapy (PICI) scholar. DM was supported through The American Association of Immunologists Intersect Fellowship Program for Computational Scientists and Immunologists.

## Author Contributions

JC-B and EJW conceived and designed the experiments. JC-B performed the experiments with help from MSA-H, YM, VC, DM, JB and MK. JC-B prepared libraries and performed sequencing with help from DM and ACH. JC-B analyzed the scRNAseq and CITE-seq datasets with help and guidance from SM. ATACseq datasets were analyzed by SM and HH in collaboration with JC-B. ZZ and SL-B provided technical help for CnR and CnR datasets were analyzed by HH in collaboration with JC-B. LS, LP and K-CG provided the orthogonal IL2Rβ construct. MK adapted the STAT5CA construct to our MSCV based expression system. JC-B and EJW wrote the manuscript with input from MSA-H, JRG, HD and ACH.

## Data availability

All sequencing data generated during this study will be made publicly available at the time of publication.

## Supplemental figure legend

**Figure S1 (related to Figure 1):**
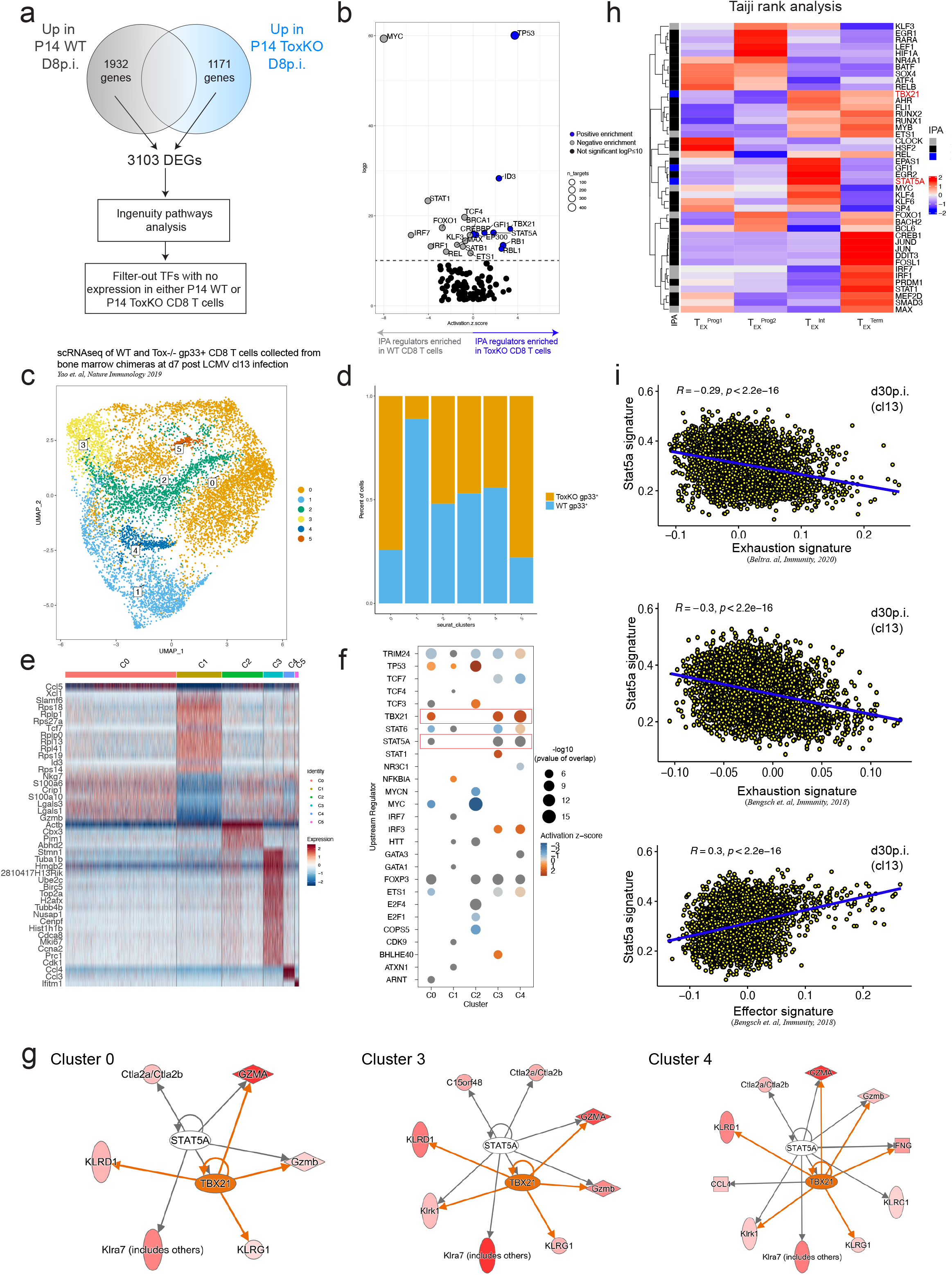
Increased Stat5a activity in T_EX_^int^ and ToxKO CD8 T cells. **A**-Analytical approach. **B**-Ingenuity <u>P</u>athways <u>A</u>nalysis on DEGs between P14 WT and P14 ToxKO cells at d8p.i. (dataset from *Khan et. al, Nature 2019*).^7^ Transcriptional regulators significantly enriched (logp>10) in P14 WT or P14 ToxKO are highlighted in grey and blue respectively. Non-significant hits are colored in black. Bubble size represents the number of genes considered by the IPA analysis for each individual TF. **C**-UMAP of re-processed scRNAseq data of WT and Tox^-/-^ gp_33_-specific CD8 T cells isolated from bone marrow chimeras at d7 post LCMV Cl13 infection and featuring Seurat clusters.^6^ **D**-Histogram showing the relative proportion of WT and Tox^-/-^ CD8 T cells in each Seurat cluster from **Fig. S1C**. **E**-Heatmap of top10 DEGs between WT and Tox^-/-^ gp33^+^ CD8 T cells per indicated cluster identified in **Fig. S1C**. **F**-IPA analysis of DEGs between WT and Tox^-/-^ gp33^+^ CD8 T cells for each individual cluster defined in **Fig. S1C**. Plotted are the top 10 transcriptional regulators enriched in each individual cluster. **G**-Network analysis of Stat5a and T-bet (*Tbx21*) by Ingenuity of DEGs between WT and Tox^-/-^ gp33^+^ CD8 T cells in indicated clusters form **Fig. S1C**. Darker red in target genes indicates positive enrichment in Tox^-/-^ gp33^+^ CD8 T cells. **H**-Taiji rank analysis identifying TFs with increased activity in previously defined subsets of T_EX_ based on published RNAseq and ATACseq data.^27^ Plotted are overlapping TFs identified in both the IPA analysis in **Fig. S1B** and the independent Taiji analysis. **I**-Correlation scores between Stat5a network and indicated gene signatures at d30 post LCMV Cl13 infection.

**Figure S2 (related to Figure 2):**
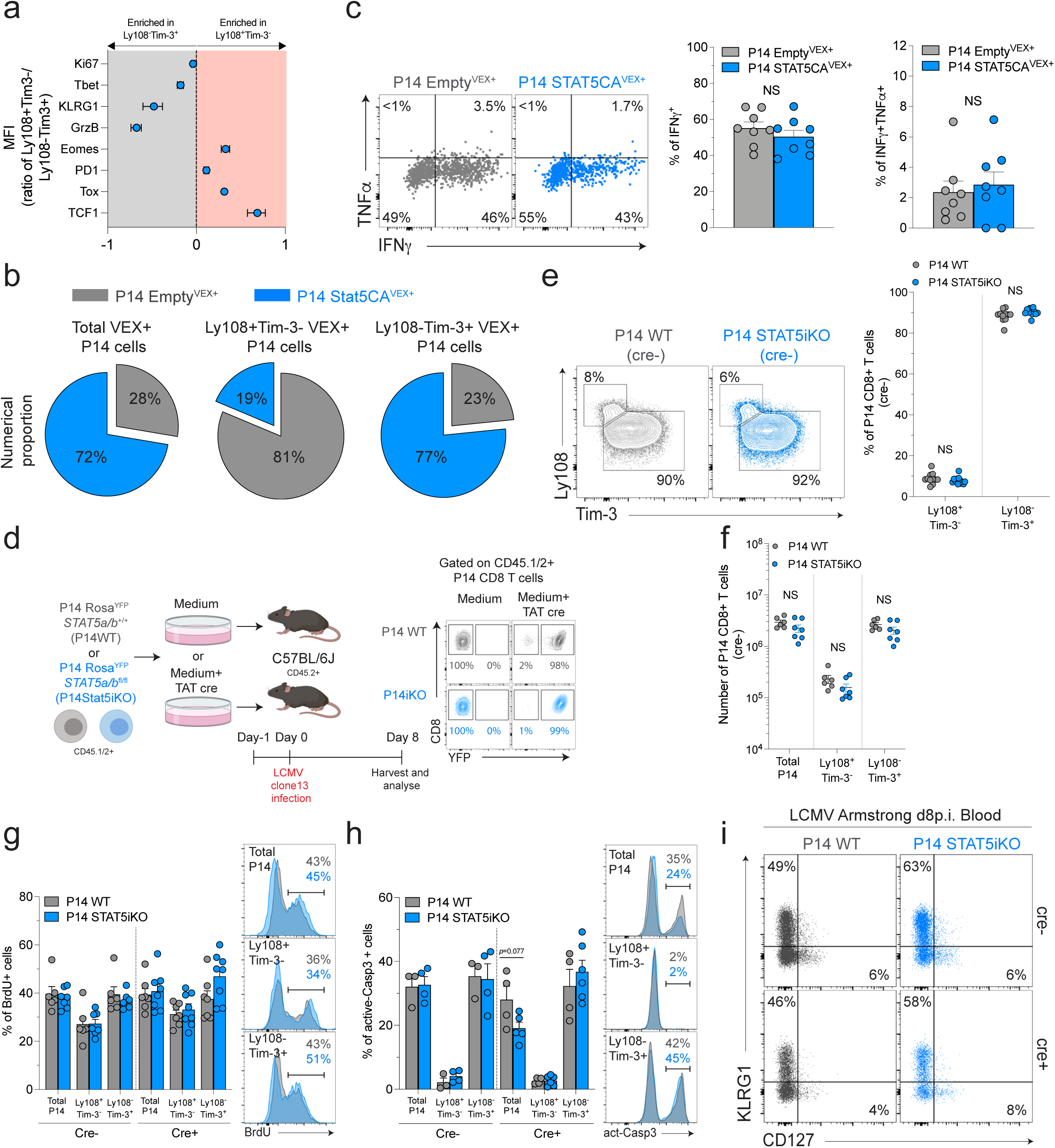
Stat5 impacts early cell-fate decision of Ag-specific CD8 T cells during a chronic viral infection. **A-** MFI of indicated markers expressed as a mean of ratio (P14 Ly108^+^Tim-3^-^/P14 Ly108^-^Tim-3^+^). N=10 with 4-20 mice/group. **B**-Numerical proportion of P14 Empty (grey) and P14 STAT5CA (blue) within indicated populations of VEX^+^ cells at d8p.i. N=4 with 16 mice/group. **C**-Representative dot plot (left) and cumulative frequencies (right) of IFNγ and TNFα production by P14 Empty (grey) and P14 STAT5CA (blue) cells at d8p.i. N=2 with 8 mice/group. **D**-Experimental design. PBMCs from CD45.1.2^+^ P14 Rosa^YFP^*Stat5a/b*^+/+^ (P14 WT) and P14 Rosa^YFP^*Stat5a/b*^fl/fl^ (P14 Stat5iKO) mice were treated (cre+) or not (cre-) with Tat cre recombinase *in vitro* and adoptively transferred into separate groups of C57BL/6J recipients subsequently infected with LCMV Cl13. Transferred cells were tracked at d8p.i. using congenic markers and YFP induction was used as a surrogate of cre-mediated recombination and deletion of floxed alleles. **E**-Representative contour plots (left) and cumulative frequencies (right) of Ly108 and Tim-3-defined subpopulations among untreated (cre-, YFP^-^) P14 WT (grey) and P14 Stat5iKO (blue) control cells at d8p.i. Numbers indicate frequencies. N=3 with 11-12 mice/group. **F**-Absolute numbers of indicated populations among untreated (cre-, YFP^-^) P14 WT (grey) and P14 Stat5iKO (blue) control cells at d8p.i. N=2 with 6-7 mice/group. **G**-*In vivo* BrdU incorporation (d7 to d8p.i.) in indicated populations of untreated (cre-, YFP^-^) and treated (cre+, YFP^+^) P14 WT (grey) and P14 Stat5iKO (blue) cells at d8p.i. **H**-Active caspase 3 staining in indicated populations of untreated (cre-, YFP^-^) and treated (cre+, YFP^+^) P14 WT (grey) and P14 Stat5iKO (blue) cells at d8p.i, after 5h incubation at 37°C. **I**-Representative dot plots of KLRG1 and CD127-defined subpopulations among treated (cre+, YFP^+^) and untreated (cre-, YFP^-^) P14 WT (grey) and P14 Stat5iKO (blue) cells isolated from the blood at d8 post LCMV Arm infection. Numbers indicate frequencies. (G-I) N=2 with 3-8 mice/group.

**Figure S3 (related to Figure 3):**
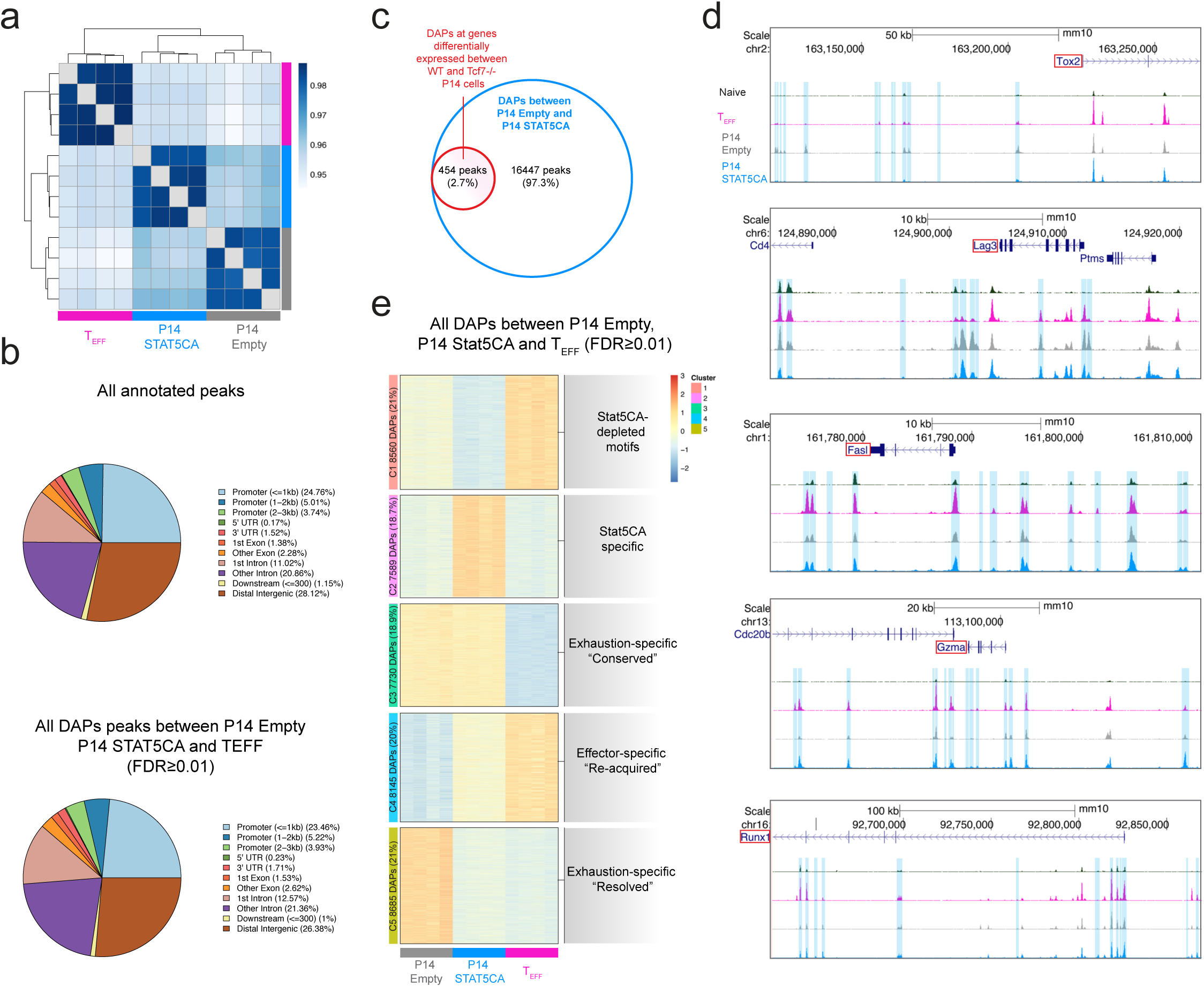
Constitutive Stat5a activity drives a hybrid epigenetic state in Ag-specific CD8 T cells. **A-** Spearman distance analysis using all DAPs (FDR≤0.01, lfc≥2) between indicated populations. Color indicates distances. **B**-Genetic distribution of the total peak list (upper panel) and all DAPs between T_EFF_, P14 Empty and P14 STAT5CA cells (lower panel; FDR≤0.01, lfc≥2). **C**-Overlap between genes containing DAPs between P14 Empty and P14 STAT5CA and genes differentially expressed between WT and *Tcf7*^-/-^ P14 cells (dataset from *Wu et. al*^33^) **D**-ATACseq tracks of indicated genes. DAPs between P14 Empty and P14 STAT5CA are highlighted in blue. **E**-Clustered heatmap (k-means) plotting all DAPs between P14 Empty, P14 STAT5CA and T_EFF_ cells (FDR≤ 0.01, lfc≥2).

**Figure S4 (related to Figure 4):**
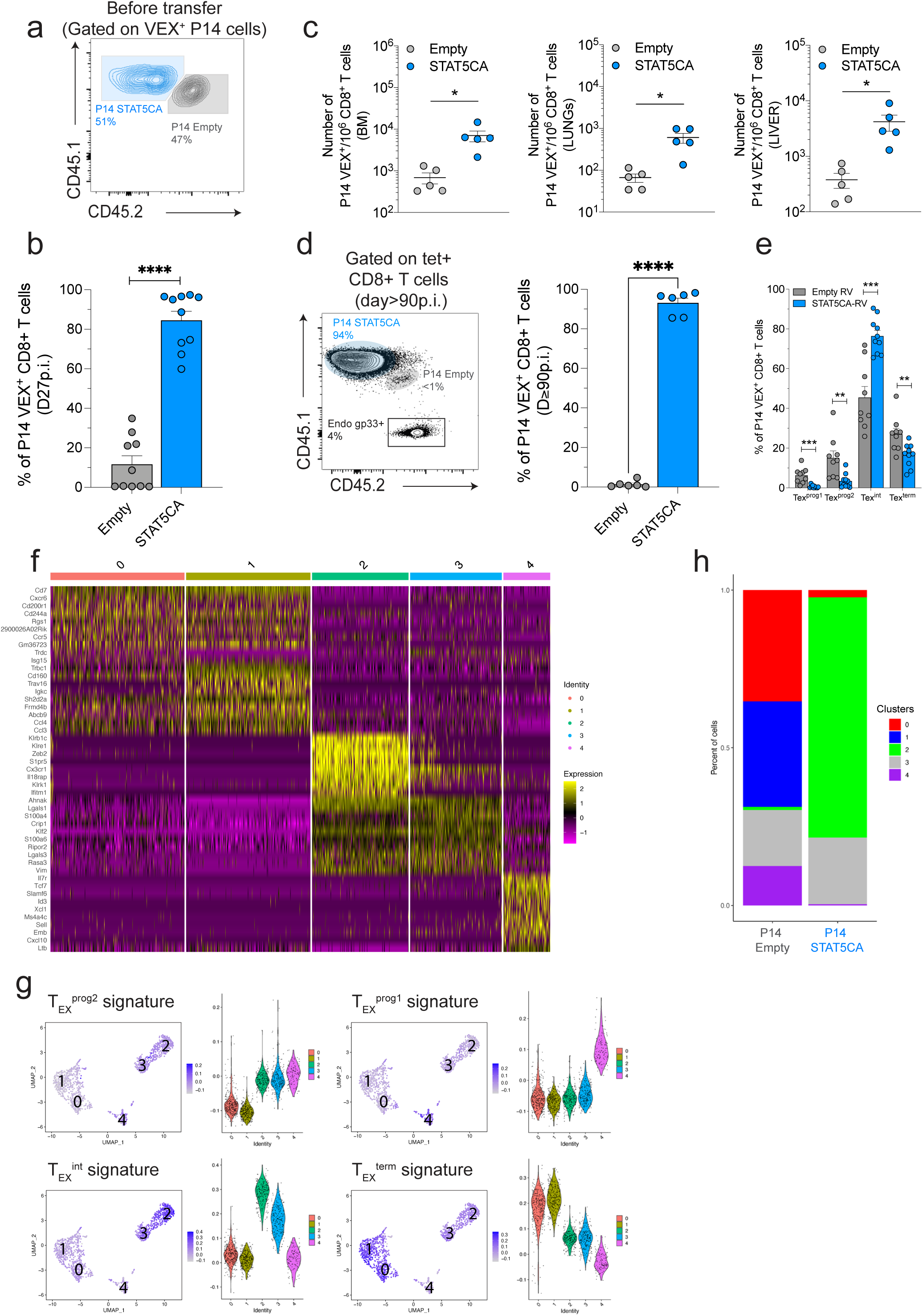
Constitutive Stat5a activity increases durability and effector biology in CD8 T cells during chronic viral infection. **A-** Pre-adoptive transfer mix plotting the relative frequency of P14 Empty^VEX+^ over P14 STAT5CA^VEX+^ prior to adoptive transfer. B-Relative frequencies of P14 Empty (grey) and P14 STAT5CA (blue) among VEX^+^ CD8 T cells at d27p.i. N=2 with 10 mice/group. C-Number of P14 Empty^VEX+^ (grey) and P14 STAT5CA^VEX+^ (blue) per 10^6^ CD8 T cells in indicated anatomical locations. Data representative of 2 independent experiments with 5 mice/group in each. D-Representative dot plot (left) and cumulative frequencies (right) of P14 Empty (grey) and P14 STAT5CA (blue) isolated from the spleen of LCMV Cl13 and CD4-depleted mice at d≥90p.i. Numbers indicate frequencies. N=2 with 6 mice/group. E-Frequencies of indicated populations among P14 Empty^VEX+^ (grey) and P14 STAT5CA^VEX+^ (blue) cells at d27p.i. N=2 with 9-10 mice/group. F-Heatmap displaying the top 10 variable genes per Seurat clusters (defined in Fig. 4D) at d27p.i. G-Projection of indicated T_EX_ subset signatures (dataset from *Beltra et. al* ^27^) into UMAP space from Fig. 4D. H-Relative distribution of indicated P14 populations across Seurat clusters (defined in Fig. 4D).

**Figure S5 (related to Figure 5):**
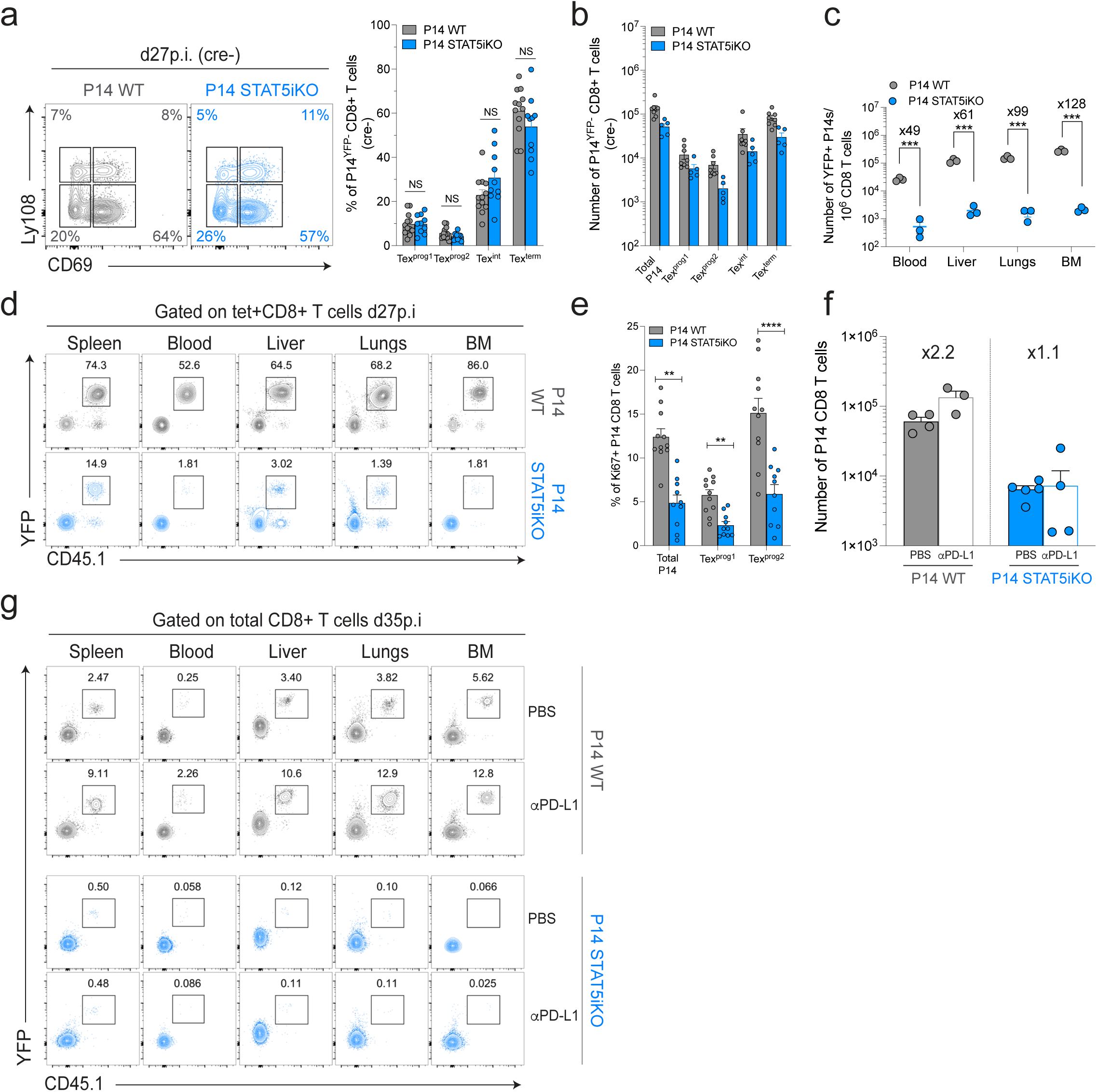
Stat5 is essential to maintain CD8 T cell responsiveness to PD-L1 blockade in settings of chronic viral infection. **A-** Representative contour plots (left) and cumulative frequencies (right) of Ly108 and CD69-defined subpopulations among untreated (cre-, YFP^-^) P14 WT (grey) and P14 Stat5iKO (blue) control cells at d27p.i. Numbers indicate frequencies. N=4 with 10-12 mice/group. B-Absolute numbers of indicated populations among untreated (cre-, YFP^-^) P14 WT (grey) and P14 Stat5iKO (blue) control cells at d27p.i. N=3 with 5-8 mice/group. C-Number of treated (cre+, YFP^+^) P14 WT (grey) and P14 Stat5iKO (blue) cells per 10^6^ CD8 T cells in indicated anatomical location at d27p.i. Numbers indicate fold changes in cell numbers. D-Representative dot plots displaying the frequency of treated (cre+, YFP^+^) P14 WT (grey) and P14 Stat5iKO (blue) cells among gp33^+^ CD8 T cells in indicated anatomical locations at d27p.i. (C-D) N=1 with 3 mice/group. E-Cumulative frequencies of Ki67^+^ cells among indicated populations of P14 WT and P14 Stat5iKO cells at d27p.i. N=3 with 10-11 mice/group. F-Absolute numbers of P14 WT (grey) and P14 Stat5iKO (blue) cells at d35p.i. in the spleen of LCMV Cl13 infected mice (depleted of CD4 T cells) and treated with either PBS or αPD-L1 blocking antibodies. Numbers indicate fold changes in cell number. G-Representative dot plots displaying the frequency of treated (cre+, YFP^+^) P14 WT (grey, upper panel) and P14 Stat5iKO (blue, lower panel) cells among CD8 T cells in indicated anatomical locations and experimental group at d35p.i. (F-G) Representative of 2 independent experiments with 3-5 mice/group in each.

**Figure S6 (related to Figure 5):**
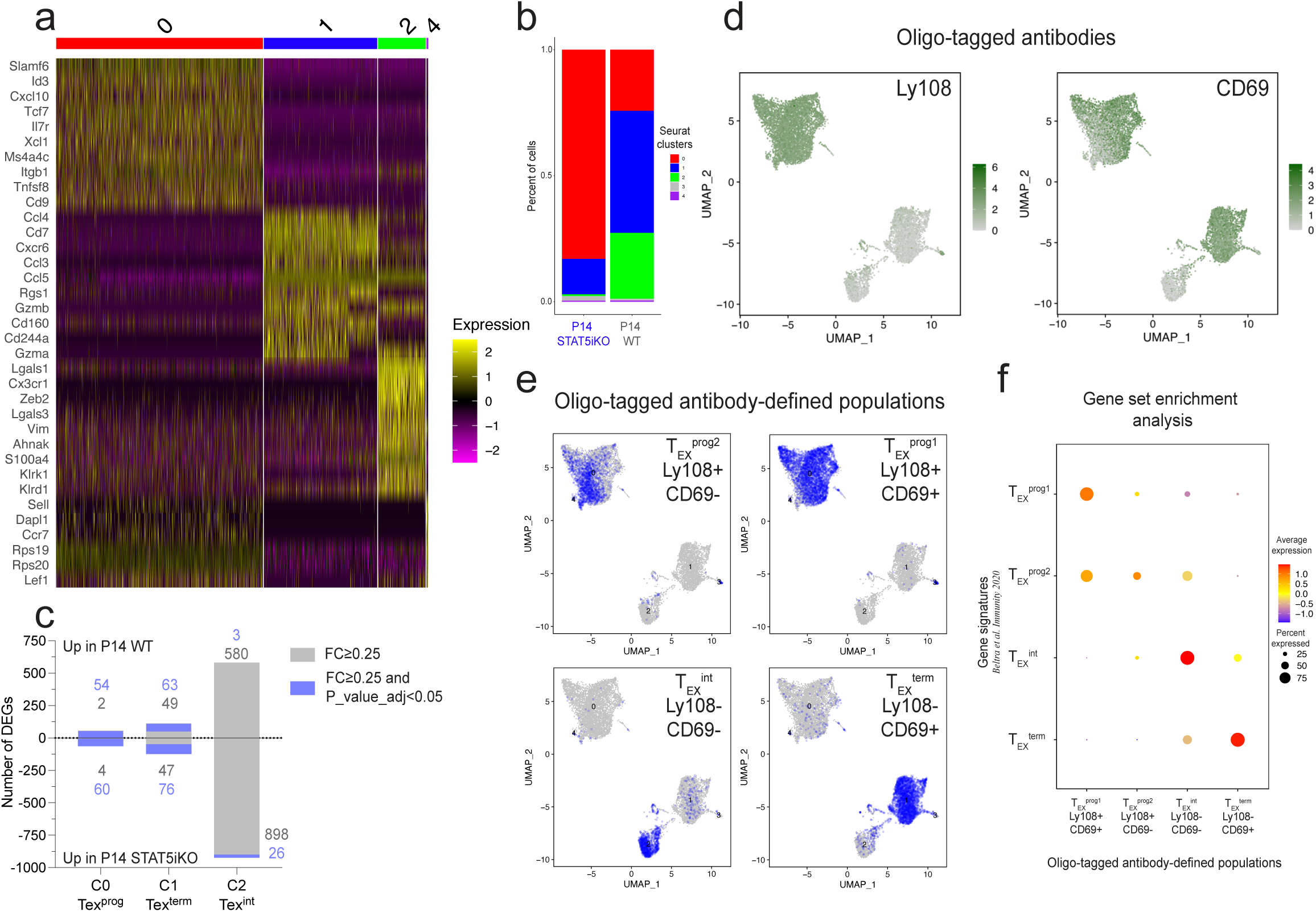
CITE-seq-mediated delineation of T_EX_ subsets. **A-** Heatmap displaying the top representative genes per Seurat clusters (defined in Fig. 5F) at d27p.i. **B-** Relative distribution of indicated populations across Seurat clusters (defined in Fig. 5F). **C**-Number of DEGs between P14WT and P14 Stat5iKO cells in indicated mRNA-based Seurat clusters. Numbers in grey indicate DEGs with lfc≥0.25 and numbers in blue DEGs with lfc≥0.25 and P_value_adj<0.05. **D**-Detection of oligo-tagged antibodies against Ly108 and CD69 across mRNA-defined Seurat clusters (defined in Fig. 5F)**. E**-Projection of indicated oligo-tagged antibodies-defined populations within mRNA-defined Seurat clusters (defined in Fig. 5F)**. F-** GSEA for indicated T_EX_ subsets signature (dataset from *Beltra et. al* ^27^) across indicated oligo-tagged antibodies-defined populations in P14 WT cells.

**Figure S7 (related to Figure 6):**
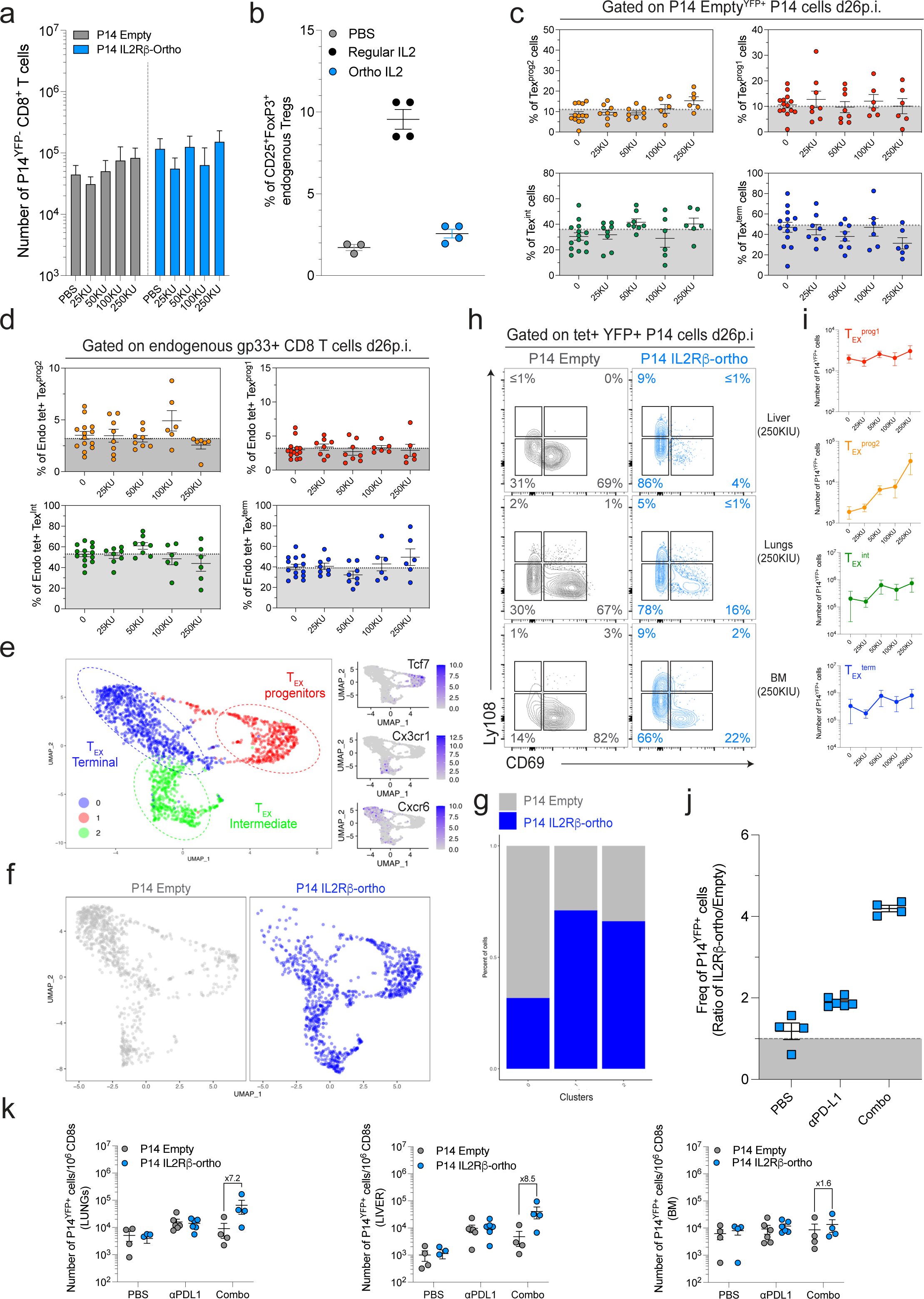
Targeted delivery of IL-2/Stat5 signals using an orthogonal IL-2/IL2Rβ pair system alters T_EX_ subset dynamic at steady state and upon PD-L1 blockade. **A-** Absolute numbers of splenic P14 Empty^YFP-^ (grey) and P14 IL2Rβ-ortho^YFP-^ (blue) cells isolated at d26p.i. from experimental groups treated with indicated doses of *ortho*IL-2 between d21-25p.i. N=2 with 6-14 mice/group. B-Frequency of endogenous Tregs (CD4^+^CD25^+^FoxP3^+^) in groups of LCMV Cl13 mice treated with PBS (grey), regular IL2 (mrIL2, 25KIU,black) or *ortho*IL-2 (25KIU, blue) between d21 and 25p.i. Data were collected from the blood at d26p.i.. N=1 with 3-4 mice/group. C-Cumulative frequencies of indicated sub-populations among P14 Empty^YFP+^ cells isolated at d26p.i. from experimental groups infused with indicated doses of *ortho*IL-2. Dotted grey lines indicate mean frequencies of each sub-population across all experimental groups in P14 Empty^YFP-^control cells. D-Cumulative frequencies of indicated sub-populations among endogenous gp33^+^ CD8 T cells isolated at d26p.i. from experimental groups infused with indicated concentrations of *ortho*IL-2. Dotted grey lines indicate mean frequencies of each sub-population across all experimental groups. (C-D) N=2 with 6-14 mice/group. E-UMAP of scRNAseq data combining P14 Empty^YFP+^ and P14 IL2Rβ-ortho^YFP+^ cells isolated at d26p.i. plotting Seurat clusters (E-left) or highlighting individual samples (F) and expression of representative genes per cluster (E-right). G-Relative distribution of indicated populations across Seurat clusters (defined in Fig. S7E). H-Representative contour plots of Ly108 and CD69 expression in co-transferred populations of P14 Empty^YFP+^ (grey) and P14 IL2Rβ-ortho^YFP+^ (blue) cells in indicated organs from mice treated with 250KIU of *ortho*IL-2 at d26p.i. Representative of 2 independent experiments with 2-6 mice/group. I-Absolute numbers of indicated populations among P14 IL2Rβ-ortho^YFP+^ cells isolated at d26p.i. from indicated experimental groups. N=2-3 independent experiments with 6-11 mice/group J-Frequency of YFP^+^ cells expressed as a ratio of splenic P14 IL2Rβ-ortho/P14 Empty at d26p.i. in indicated experimental groups. Representative of 2 independent experiments with 9-11 mice/group. K-Number of P14 Empty^YFP+^ (grey) and P14 IL2Rβ-ortho^YFP+^ (blue) per 10^6^ CD8 T cells in indicated anatomical locations and experimental groups. N=1 with 3-6 mice/group.

**Figure S8 (related to Figure 7):**
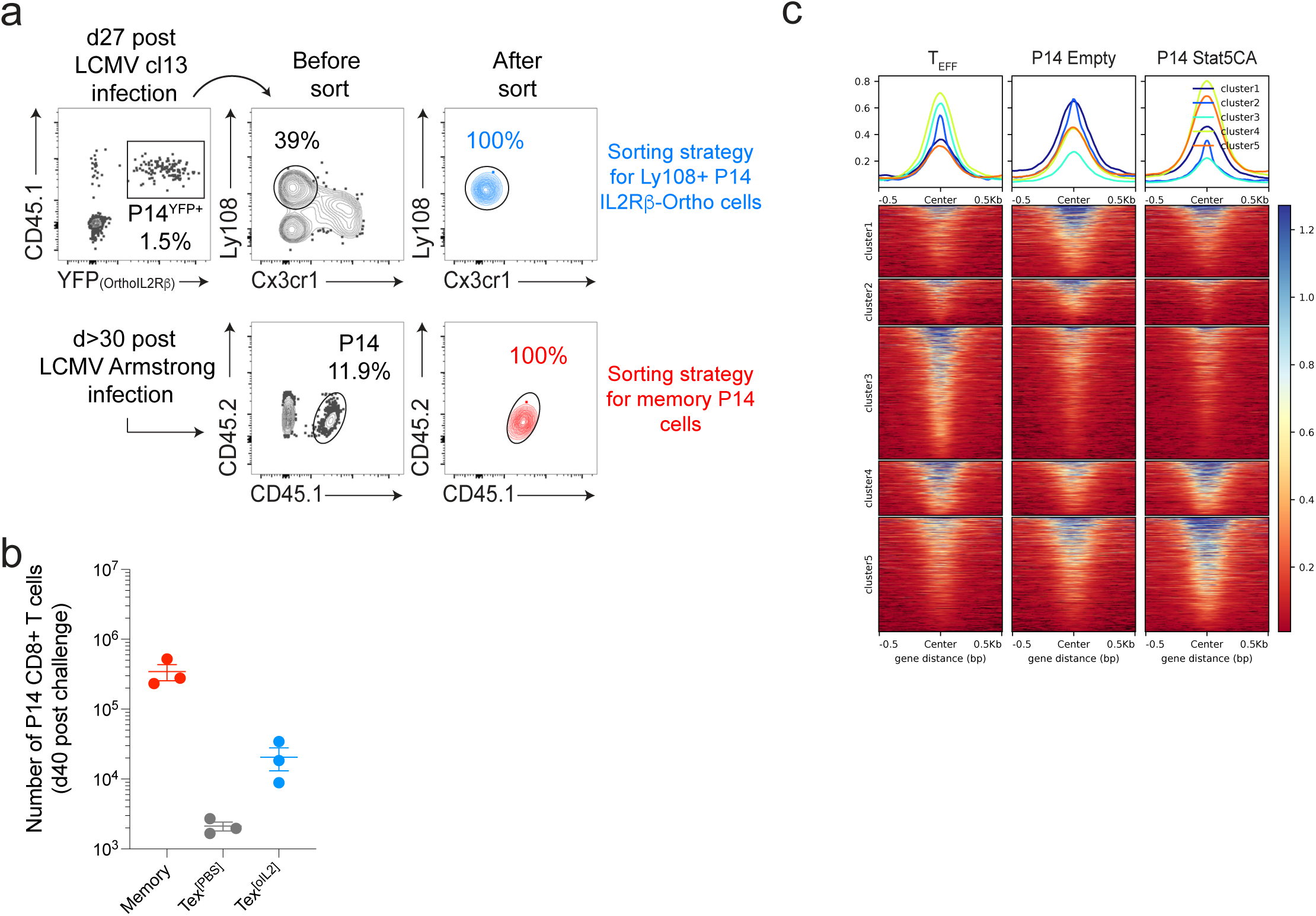
Targeted delivery of IL-2/Stat5 signals during viral rechallenge alters T_EX_ biology. **A-** Sorting strategy for P14 T_MEM_ (lower panel) and P14 IL2Rβ-ortho cells (Ly108^+^ T_EX_ progenitors expressing the IL2Rβ-ortho receptor [YFP reporter], upper panel). **B**-Absolute numbers of indicated P14 cell populations in the spleen at d40 post re-challenge. N=1 with 3 mice/group. **C**-Heatmap for scores associated with genomic regions plotting accessibility at genomic regions identified in Fig. 7H for T_EFF_, P14 Empty and P14 STAT5CA cells isolated at d8 of Cl13 infection as detailed in Fig.3.

## Tables

Table S1: IPA results and gene input

Table S2: DAPs between P14 Empty, P14 STAT5CA and T_EFF_

Table S3: DEGs between C2 and C3 (Fig. 4G)

Table S4: Genes with increased accessibility in P14 STAT5CA vs P14 Empty at d8p.i.

Table S5: DEGs between P14WT and P14 Stat5iKO

Table S6: DAPS between rechallenged Memory, T_EX_^[PBS]^ and T_EX_^[oIL2]^

